# *MLC1* alteration in iPSCs give rise to disease-like cellular vacuolation phenotype in the astrocyte lineage

**DOI:** 10.1101/2025.01.06.631607

**Authors:** Saumya Sharma, Vishal Bharti, Prosad Kumar Das, Abdul Rahman, Harshita Sharma, Riya Rauthan, Madhumita RC, Neerja Gupta, Rashmi Shukla, Sujata Mohanty, Madhulika Kabra, Kevin R Francis, Debojyoti Chakraborty

## Abstract

**Background:** Megalencephalic leukoencephalopathy with subcortical cysts (MLC), a rare and progressive neurodegenerative disorder involving the white matter, is not adequately recapitulated by current disease models. Somatic cell reprogramming, along with advancements in genome engineering, may allow the establishment of *in-vitro* human models of MLC for disease modeling and drug screening. In this study, we utilized cellular reprogramming and gene-editing techniques to develop induced pluripotent stem cell (iPSC) models of MLC to recapitulate the cellular context of the classical MLC-impacted nervous system.

**Methods:** Somatic cell reprogramming of peripheral patient-derived blood mononuclear cells (PBMCs) was used to develop iPSC models of MLC. CRISPR-Cas9 system-based genome engineering was also utilized to create the *MLC1* knockout model of the disease. Directed differentiation of iPSCs to neural stem cells (NSCs) and astrocytes was performed in a 2D cell culture format, followed by various cellular and molecular biology approaches, to characterize the disease model.

**Results:** MLC iPSCs established by somatic cell reprogramming and genome engineering were well characterized for pluripotency. iPSCs were subsequently differentiated to disease-relevant cell types: neural stem cells (NSCs) and astrocytes. RNA sequencing profiling of MLC NSCs revealed a set of differentially expressed genes related to neurological disorders and epilepsy, a common clinical finding within MLC disease. This gene set can serve as a target for drug screening for the development of a potential therapeutic for this disease. Upon differentiation to the more disease relevant cell type-astrocytes, MLC-characteristic vacuoles were clearly observed, which were distinctly absent from controls. This emergence recapitulated a distinguishing phenotypic marker of the disease.

**Conclusion:** Through the creation and analyses of iPSC models of MLC, our work addresses a critical need for relevant cellular models of MLC for use in both disease modeling and drug screening assays. Further investigation can utilize MLC iPSC models, as well as generated transcriptomic data sets and analyses, to identify potential therapeutic interventions for this debilitating disease.

## INTRODUCTION

Megalencephalic Leukoencephalopathy with subcortical cysts (MLC) is a slowly progressive degenerative brain disease involving the white matter which is a result of a spectrum of pathogenic variants across *MLC1* or *GlialCAM* genes. This disease was first discovered independently by Dr. Marjo Van der Knaap in the Netherlands (van der Knaap et al., 1995) and by Dr. Bhim Sen Singhal in the Indian Agrawal community (Singhal et al., 1996). Therefore, MLC is also named as Van der Knaap-Singhal disease (van der Knaap et al., 2012). The three major categories of causal variations are: Autosomal recessive mutation in *MLC1*, an autosomal recessive and an autosomal dominant mutation in *GlialCAM* (Capdevila-Nortes et al., 2013). *MLC1* was the first gene known to cause MLC and was mapped to chromosome 22qtel (Topçu et al., 2000; Leegwater et al., 2001). *MLC1* translates into a protein (MLC1) that is mainly expressed in astrocytes within the brain, especially at astrocytic end feet that contact the blood-brain barrier (Masaki et al., 2012), within the pia mater, and within astrocytes present at the synaptic cleft (Kater et al., 2023). The structural features and observed brain defects in patients of MLC, such as brain oedema, fluid filled cysts, vacuolation in astrocytes and hypomyelination, suggest that MLC1 may regulate ionic and water balance within the brain (Ridder et al., 2011; Lanciotti et al., 2012; Brignone et al., 2014; Hoegg-Beiler et al., 2014; Dubey et al., 2014). A fully elucidated role of MLC1 protein is yet to be deciphered. In MLC, patients develop macrocephaly from an early age with other characteristics such as developmental delay, intellectual disability, motor deficits, seizures, mental decline, and ataxia. Though severity is variable, it is usually a slowly progressive disease and individuals may live into their fifties (Knaap et al., 2018). MLC is found prevalent in the Turkish (van der Knaap et al., 1995) (Yiş et al., 2009) and Indian Aggarwal (Singhal et al., 1996; Singhal et al., 2003; Gorospe et al., 2004; Singhal, 2005) communities with some cases found in Japan (Saijo et al., 2003), Israeli community (Ben-Zeev et al., 2001), Finland (Leegwater et al., 2001), Italy (Montagna et al., 2006), Egypt (Abdel-Salam et al., 2016) and Korea (Choi et al., 2017) as well.

At present, a cure for MLC is not available and patients rely on supportive treatments. Current management of MLC disease is essentially based on the specific set of symptoms observed in the patients and requires execution of multidisciplinary approaches working towards the goal of seizure control, physical therapy, and avoiding head trauma to the patient (which can temporarily worsen the symptoms) (Batla A et al., 2011. These management strategies only work as palliative cure and require an absolute therapeutic solution for MLC disease in its repertoire. Since it is a rare genetic brain disease with approximately 200 reported cases worldwide, research towards discovering a therapeutic intervention is needed. Thus, a relevant disease model is needed to identify potential therapies for this debilitating disease.

To understand MLC pathology, various *in-vitro* and *in-vivo* models have been developed. Expressing mutant MLC1 protein in heterologous cell lines (such as HeLa, HEK293 cells and Xenopus oocytes) indicated that missense variations in *MLC1* gene are more often pathological. They hinder plasma membrane localization of MLC1 protein (Teijido et al., 2004). Some of these variations also affect the half-life of MLC1 protein by resulting in an increased degradation of this protein: misfolding of protein resulting in an activation of the endoplasmic reticulum associated degradation (ERAD) pathway (Duarri et al., 2008). MLC1 protein is highly expressed in astrocytes; therefore, astrocytes represent a relevant *in-vitro* model system to study MLC1 pathological effects (Ambrosini et al., 2008). The human astrocytoma cell line U251 cells, that expresses almost undetectable levels of endogenous MLC1 protein, were made to stably express *MLC1* gene or *MLC1* gene with several missense/ pathological variations. It was discovered that not every mutant protein is able to reach the plasma membrane and might be retained in the endoplasmic reticulum (ER) (Brignone et al., 2010). Many pathogenic variations altered interaction of MLC1 with its bonafide interactors, such as GlialCAM, Kir4.1, Na K-ATPase, V-ATPase, and TRPV4 (Lanciotti et al., 2012). This showed that *in-vitro* modeling of *MLC1* patient variations could specifically reveal variation-specific dysfunctionality and these models can then be used to develop patient-specific therapeutics. Mlc1 downregulation in primary rat astrocytes, with small interfering RNA (siRNA) and *Mlc1* deletion in mice (Dubey et al., 2014) resulted in emergence of intracellular vacuoles, a pathological hallmark of the MLC affected brain (Duarri et al., 2011). However, this system did not recapitulate the mis-localization event of MLC1 interacting protein ZO-1 as seen in patients. Other mouse models have been generated with complete knockdown of Mlc1 across all tissues (Hoegg-Beiler et al., 2014; Dubey et al., 2014). *Mlc1* mouse models recapitulate some aspects of human MLC: early onset of disease, increased brain water content, cysts in the white matter, and abnormal astrocytes near the blood brain barrier. Additionally, a zebrafish model has also been developed by deleting the *MLC1* ortholog (Sirisi et al., 2014). However, the severity of the disease condition in all of these models do not fully recapitulate the actual human condition, with the zebrafish model bearing even lesser severity of the disease-like condition. Poor disease recapitulation may be due to differences between species or due to unknown compensatory mechanisms in these model organisms that are possibly absent from the human subjects. Besides, in higher mammals, the glia population is known to be higher in complexity and the glia to neuron ratio is also significantly higher (Pfrieger, 2009). These findings limit the utility of studying MLC in currently available animal models.

With advances in somatic cell reprogramming (Takahashi & Yamanaka, 2006) and CRISPR-Cas9 genome editing (Gasiunas et al., 2012)(Jinek et al., 2012), multiple new avenues were opened for rare disease research. Neural differentiation and patterning of derived induced pluripotent stem cells (iPSC) can model a diseased brain in a dish in an indirect manner, without causing harm to the donor (Vadodaria et al., 2018). Temporal exposure of iPSCs to important signaling molecules has given rise to protocols that can be used to generate distinct populations of neurons and glia from iPSCs (Liu & Zhang, 2011), including astrocytes. Astrocytes are the primary cell type within the mammalian brain (Shao & McCarthy, 1994). Astrocytes help neurons by guiding their development and parallelly support them in terms of nutrition and metabolism (Rudge, 1993). They have also been associated with multiple brain pathologies, where defects in their functionality can impact neuronal survival and neurologic function (van der Laan et al., 1997). Astrocytes express certain surface molecules and release factors that affect recovery of the brain after an injury, outgrowth of neurites, synaptic plasticity or neuron regeneration (Chen & Swanson, 2003). To understand MLC, which has a direct link with astrocyte activity, it is essential to analyze MLC impacts on disease-relevant neural cell types such as astrocytes to identify disrupted signaling pathways and cellular deficits within MLC.

## METHODS

### PBMC isolation and reprogramming to iPSCs

10 ml of freshly drawn blood was diluted with PBS (1:1) and the dilution is layered over Ficoll Paque (GE 17- 1440-02) (Day -4). This is followed by centrifugation (330 g, 45 min at 21°C, 1 unit acceleration and 0 unit deceleration) based separation, to obtain peripheral blood mononuclear cells (PBMCs) at the “buffy coat”. Extracted out the contents of the buffy coat and washed them twice in 5% FBS + 1x DPBS (330 g, 15-30 min at 21°C, 9 units acceleration and deceleration). Part of the freshly isolated PBMCs were frozen down and the other part, cultured in PBMC media [StemPro®-34 SFM Complete Medium (Gibco™) supplemented with L-Glutamine (2mM)] with cytokines [FLT3 (100 ng/ml; Cat. No.: PHC9415), SCF (100 ng/ml; Cat. No.: PHC2116), IL-3 (20 ng/ ml; Cat. No.: PHC0034), and IL-6 (20 ng/ml; Cat. No.: PHC0065)] for 4 days with daily, half media changes (designated day -4, day -3, day -2 & day -1). On day 0, for reprogramming PBMCs to iPSCs, cells were collected by centrifugation (200 g, 5 min at 21°C) and treated with Sendai virus-based vectors for transduction (Fusaki et al., 2009), as per the manufacturer’s instructions given for CytoTune-iPSC 2.0 Sendai Reprogramming kit. On the next day (day 1), cells were collected, centrifuged, and then rinsed with PBMC media to remove viral vectors from spent media. Cells were transferred to an ultra-low attachment plate and incubated for two days without any media changes. On day 3, cells were collected and plated on a fresh, Matrigel-coated, 6 well plate (20,000 - 50,000 cells per well) in PBMC media without cytokines and incubated without disturbing until day 6. On day 6, the entire spent PBMC media was replaced with fresh PBMC media. Transitioning to mTesr1 media began with half media replacement from day 7, followed by a complete media replacement and replenishment from day 8 to day 21 while observing cells for morphological changes. After day 21, ESC-like cells were isolated by manual picking of live stained Tra-1-60 positive colonies and transferring these individually to a Matrigel-coated, 24 well plate. These harvested colonies were qualified to be prospective iPSC clones, subjected to further characterizations, allowing them to amplify and propagate onto Matrigel-coated plates and passaging (at the ratio of 1:4) with ReLeSR ( **Table S3**).

### Generation of *MLC1* iPSC models with CRISPR-Cas9 knockout

A single cell suspension was made of NL5 iPSCs (NCRM-5; a kind gift from the National Heart, Lung, and Blood Institute iPSC core facility, NIH, Bethesda, MD), with the help of accutase. 300,000 cells were counted and pelleted. For each cell pellet, pSpCas9(BB)-2A-GFP (PX458; a kind gift from Feng Zhang, Addgene plasmid #48138) with two guide RNAs targeting exon 1 and exon 2 of *MLC1* gene was taken and a cell + plasmid DNA suspension was made in 15 μl of Buffer R (Neon Electroporation system, Invitrogen). Parallely, the same dual guide plasmid but with scrambled sequence of crRNA was used as control. Electroporation was carried out by dipping the gold coated tip carrying 10 ul of the suspension into 3 ml of Buffer E (Neon electroporation system, Invitrogen); electroporation parameters were: 1200 V, 30 ms, 1 pulse. The tip contents were thereafter released into a single well of a matrigel coated 24 well plate, now containing STEMacs iPS Brew (Miltenyi Biotec) plus CloneR (Stem Cell Technologies) reagent. The cells were then allowed to repair and replenish for the next 24 hrs, after which a media change was performed. 48 hrs post-electroporation, the cells were collected and a flow cytometry based single cell sorting was performed into a 96 well plate, coated with Matrigel, and containing stem cell media plus CloneR. Sorted cells were allowed to grow until proper colonies were visible and individual cell clones could thereafter be propagated, cryostored, and characterized for pluripotency.

### Characterization of reprogrammed iPSCs and *MLC1* knockout iPSCs

#### Tra-1-60 immunocytochemistry

Cells were collected in PBS with 5% BSA. DyLight 488-conjugated Tra-1-60 antibody [(1:100); Cat no.: MA1-023-D488X; Invitrogen] was added to cells, incubated on ice for 45 mins, washed 3x with PBS, strained through a 40 μm membrane, and analyzed by FACS to assess efficiency of pluripotency attainment.

#### For pluripotency determination

iPSC clones were subjected to alkaline phosphatase assay as per manufacturer protocol. Patient derived iPSCs were stained with Tra-1-60 antibody. A panel of antibodies were used to assess pluripotency (**Table S3**).

#### Trilineage differentiation of iPSCs

Patient-derived iPSCs were subjected to trilineage differentiation using StemDiff trilineage differentiation kit [Cat no.: 05231-Ecto; 05232-Meso; 05233-Endo; Stem Cell Technologies] following manufacturer’s instructions. For MLC1 knockout iPSCs and scrambled controls, an undirected differentiation protocol was used. Embryoid bodies (EBs) were prepared by forced aggregation of 200-250 cells in an Aggrewell 800 plate in iPSC media without bFGF for 6 days. After 6 days, the EBs were plated to 0.1% gelatin-covered dishes in DMEM-F12 media with 20% FBS for 14 days. Trilineage marker profiling was done by performing a qPCR assay with TaqMan probe based qPCR conditions (Primer & oligo list in **Table S1** & **Table S2**).

#### Immunofluorescence (IF) staining

The expression of pluripotency and tri-lineage markers was studied using IF staining. Cells were fixed in 4% paraformaldehyde (PFA) for 15 min at room temperature and after PBS washes, incubated in blocking buffer [10% normal goat serum (NGS), 1% BSA and 0.3% Triton X-100 in phosphate buffer saline (PBS)] for 1 hr at room temperature. Afterwards, cells were incubated with primary antibodies diluted in their respective dilution buffers (for trilineage differentiation assay: Sox1 (in 1% BSA, 1% NGS, 0.25% Triton-X in PBS), Alpha-SMA (in blocking buffer), GATA4 (in 1% BSA and 0.1% Tween20)] and for pluripotency assay: Oct4, Sox2 and Klf4] at 4 °C overnight and then incubated with secondary antibodies for 1 hr at room temperature. Finally, nuclei were stained with DAPI for 15 min at room temperature. Post staining, cells were visualized and imaged under a fluorescence microscope (FLoid™ Cell Imaging Station).

#### Alkaline phosphatase assay

Live cells were treated with alkaline phosphatase staining dye following manufacturer protocols. Purple/red color is an indicator of pluripotency in the iPSC colony.

#### Assessment of karyotype

For qPCR based karyotyping, a commercially available kit (Stem Cell Technologies) was used per the manufacturer’s instructions to determine the status of eight commonly observed karyotype anomalies in reprogrammed iPSCs.

#### Mycoplasma test

Mycoplasma analysis was performed at different stages, using either of two detection methods: a Mycoplasma Detection Kit (MycoAlert, Lonza) or PCR based assessment of mycoplasma in cultured cells. The culture medium and gDNA was used as the test sample to screen for mycoplasma contamination according to manufacturer’s protocol.

#### Mutation analysis

Genomic DNA was extracted and purified from DBT-MLC2 iPSCs. The MLC region flanking the mutation site was amplified and cloned into the pcr2.1 TOPO vector. Six colonies were picked at random and the plasmid preparation was analyzed by Sanger sequencing.

### Differentiation of iPSCs to Neural Stem Cells

0.9 to 3 million cells in single cell suspension format were counted and seeded in each well of a pre-rinsed Aggrewell 800 plate per manufacturer protocol (Stem Cell Technologies) in 2 ml iPSC media with Y27632 ROCK inhibitor (10 μM, Stem Cell Technologies). The plate was centrifuged at 100 x g, 3 min, in a swinging bucket centrifuge to settle down the cells within the microwells of each well of the Aggrewell plate in a homogenous manner. Next day and onwards, generated EBs were transitioned to neural induction media: DMEM without glutamine or sodium pyruvate, N2B supplement, B27 with Vitamin A, EGF (20ng/ml), bFGF (20ng/ml), LDN193189 (Reagents Direct, 100 nM), 1x L-glutamine, SB431542 (Reagents Direct, 10 uM), and pen-strep (100 units/mL). Half media changes were performed every other day until day 8. On day 8, neuralized EBs were carefully picked up with help of a wide bore pipette tip and allowed to sediment with gravity. Sedimented EBs were resuspended in fresh media and were plated on a poly-L-ornithine (PLO) and laminin coated 60 mm dish. To coat with PLO/Laminin, the following procedure was followed: on -2 day, each dish was coated with 20 ug/ml PLO in nuclease free water (NFW). Kept the plate at 37 °C overnight. On -1 day, removed PLO coating solution, washed 2x with NFW, followed by coating with 10 ug/ml Laminin in DPBS based solution (for plastic dishes); 20 ug/ml laminin solution was used for glass dishes, 37°C incubator overnight. Freeze the plates in -20 and use them for upto 6months). Next day onwards, media was transitioned from EB media to NSC media. NSC media: DMEM (-Glutamine; -Sodium Pyruvate), 1x L-Glutamine, EGF (20 ng/ml), bFGF (20 ng/ml), B27 (- Vitamin A), 1x PenStrep. 2-3 days post EB transfer, neural rosettes were observed, but surrounded by neural crest population. Careful separation of just the neural rosettes was done by carefully plucking out the rosette centers into the media, followed by plating the rosette containing supernatant onto a fresh PLO/Laminin dish. This was considered the P0 NSC population. 2-3 days later, NSCs were accutase treated and propagated further by passaging in a 1:2 manner.

### Directed differentiation of neural stem cells to astrocytes

15,000 NSCs per cm^2^ were seeded in a T25 flask in NSC media. The next day, media was transitioned to astrocyte differentiation media consisting of astrocyte media (ScienCell Research Laboratories; Carlsbad, CA, USA; Cat. No. 1801b), astrocyte growth supplement (1x; ScienCell Research Laboratories; Cat. No. 1852), 2% Fetal Bovine Serum (FBS; ScienCell Research Laboratories; Cat. No. 0010), 50 U/ml penicillin G, 50 mg/ml streptomycin. Half medium was replaced every 3-4 days and cells were passaged every week in a 1:4 ratio on poly-D-lysine (PDL) coated dishes. After 30 days, the media was transitioned to astrocyte maturation media. Cells were then cultured for another 2-3 months depending on the experimental endpoints (Brennand KJ et. al., 2011).

### RNA isolation and sequence analysis

#### Sample Preparation and RNA Sequencing

Total RNA was isolated from knockout (KO) and scrambled (wild type) NSCs. NSCs were collected in biological duplicates and used to prepare cDNA libraries. Samples were sequenced on an Illumina NovaSeq 6000 platform, generating paired-end 150 base pair reads.

#### Quality Check and Preprocessing of RNA-Seq Data

Quality control of the raw sequencing data was performed using FastQC version 0.11.9. The raw reads were trimmed to remove adapter sequences and low-quality bases using Trimmomatic version 0.39 with default settings (Bolger et al., 2014). The trimmed reads were again analyzed with FastQC and the results were consolidated using MultiQC version 1.14 (Ewels et al., 2016).

#### Alignment and Quantification

The cleaned reads were aligned to the human reference genome GRCh38 using HISAT2 version 2.2.1 (D. Kim et al., 2015) with default parameters. The resulting Sequence Alignment/Map (SAM) files were converted to Binary Alignment/Map (BAM) files, sorted and indexed using SAM tools version 0.1.15 (Li et al., 2009). Transcript quantification was performed using feature counts, producing raw gene counts for each sample (Liao et al., 2013).

#### Differential Gene Expression Analysis

Differential gene expression analysis was performed using DESeq2 version 1.36.0 in R version 4.2.2 (Love et al., 2014) (Aslam & Imdad Ullah, 2023). Genes with an absolute log2 fold change greater than 1 and a false discovery rate (FDR) adjusted p-value less than or equal to 0.05 were considered significantly differentially expressed. The Enhanced Volcano package version 1.14.0 was used to visualize the differential gene expression results in a comprehensive volcano plot (*VolcaNoseR – a Web App for Creating, Exploring, Labeling and Sharing Volcano Plots*, n.d.).

#### Functional and Pathway Enrichment Analysis

Functional enrichment analysis was conducted on the significantly differentially expressed genes using the DAVID bioinformatics resource (Huang et al., 2008). Pathway enrichment analysis was performed using KEGG (Kanehisa, 2000), Reactome (Fabregat et al., 2015), and WikiPathways (Slenter et al., 2017) databases through DAVID. Disease enrichment analysis was performed using the DisGeNET database within DAVID. The resulting disease terms were classified into different disease categories and visualized.

#### Protein-Protein Interaction (PPI) Analysis and Disease Association

The STRING database was used to investigate protein-protein interactions among the seven genes associated with neurological disease categories (Piñero et al., 2019). A heatmap was created to represent the enrichment of these genes in epilepsy-related diseases as obtained from the DisGeNET database.

#### Drug Target Analysis and ADME Prediction

The DrugBank database was used to identify potential drugs targeting the seven selected genes (Szklarczyk et al., 2018). The drugs were filtered to only include those classified as ‘Approved’, ‘Investigational’, or ‘Experimental’. The SwissADME platform was employed to perform absorption, distribution, metabolism, and excretion (ADME) predictions on the identified drugs (Wishart et al., 2017). Investigation of drug repurposing based on synergy, ADME, and the pathways that the seven genes are involved in, are ongoing.

### Screening of *MLC1* disease-causing mutation

Genomic DNA was isolated from patient PBMCs and the patient derived iPSCs. A 271 bp amplicon of *MLC1* was amplified by PCR using the above genomic DNAs with two primer sets and amplicons were sequenced using Sanger sequencing (Primer list in **Table S1**).

## RESULTS

### MLC patient-derived PBMCs were successfully reprogrammed to iPSCs

To generate an MLC disease model with a closer link to human subjects, we focused on a *MLC1* mutation that produces a severe clinical phenotype. The 132dupC mutation in the Indian subcontinent affects members of the Indian Agrawal community, resulting in a mildly progressive form of MLC (Singhal et al., 2003; Gorospe et al., 2004).

We derived PBMCs from the MLC patient and successfully reprogrammed them to an iPSC state (**Figure 1a**). PBMCs were reprogrammed using titered Sendai viral particles bearing the Yamanaka reprogramming factors. After 25 days, morphologically pluripotent cells emerged with increased volume and surface areas of some colonies, indicating successful reprogramming (**Figure 2a**). Individual iPSC clones were subsequently picked, plated, propagated and cultured separately. Subsequently, a series of characterization experiments were performed to confirm that the resulting cells had indeed adopted an iPSC identity. Colonies exhibited robust alkaline phosphatase activity (**Figure 2b**). To verify the patient-derived iPSCs had the desired MLC1 genotype, we utilized Sanger sequencing to confirm the presence of the specific *MLC1* variation in the cells (**Figure 2c**; **Supplementary Figure S2**). We also examined the expression of a suite of pluripotency markers. These markers span a range of proteins integral to maintaining the self-renewal and an undifferentiated stem cell state (**Figure 2d**). We subsequently sought to ascertain the global genomic stability of our iPSCs. This assessment is critical, as the reprogramming procedure, while enabling somatic cells to revert to a pluripotent state, can also potentially induce chromosomal abnormalities (Liu X. et. al., 2020). We first utilized a qPCR based karyotype analysis of the iPSC clones to demonstrate a lack of karyotypic change (**Supplementary Figure S3**). We also employed high-resolution G-banding to allow a comprehensive, chromosome-by-chromosome analysis of the entire karyotype. We ascertained that our iPSCs were devoid of major chromosomal anomalies, thereby confirming suitability for further research applications (**Figure 2e**). Our subsequent objective was to evaluate the potential of our patient-derived iPSCs to differentiate into all three primary germ layers representing the earliest differentiation events of embryonic development and serve as the precursors to all the tissues within an organism. An assessment of tri-lineage differentiation in our patient-derived iPSC line is a key criterion for validating pluripotency. Following a standardized differentiation protocol, we verified successful lineage commitment by staining the cells for a set of lineage-specific markers representing the ectodermal, mesodermal, and endodermal lineages. The presence of these markers post-differentiation demonstrated their capacity to differentiate into all three germ layers, thereby further substantiating their pluripotent state (**Figure 2f**).

**Figure 1.**
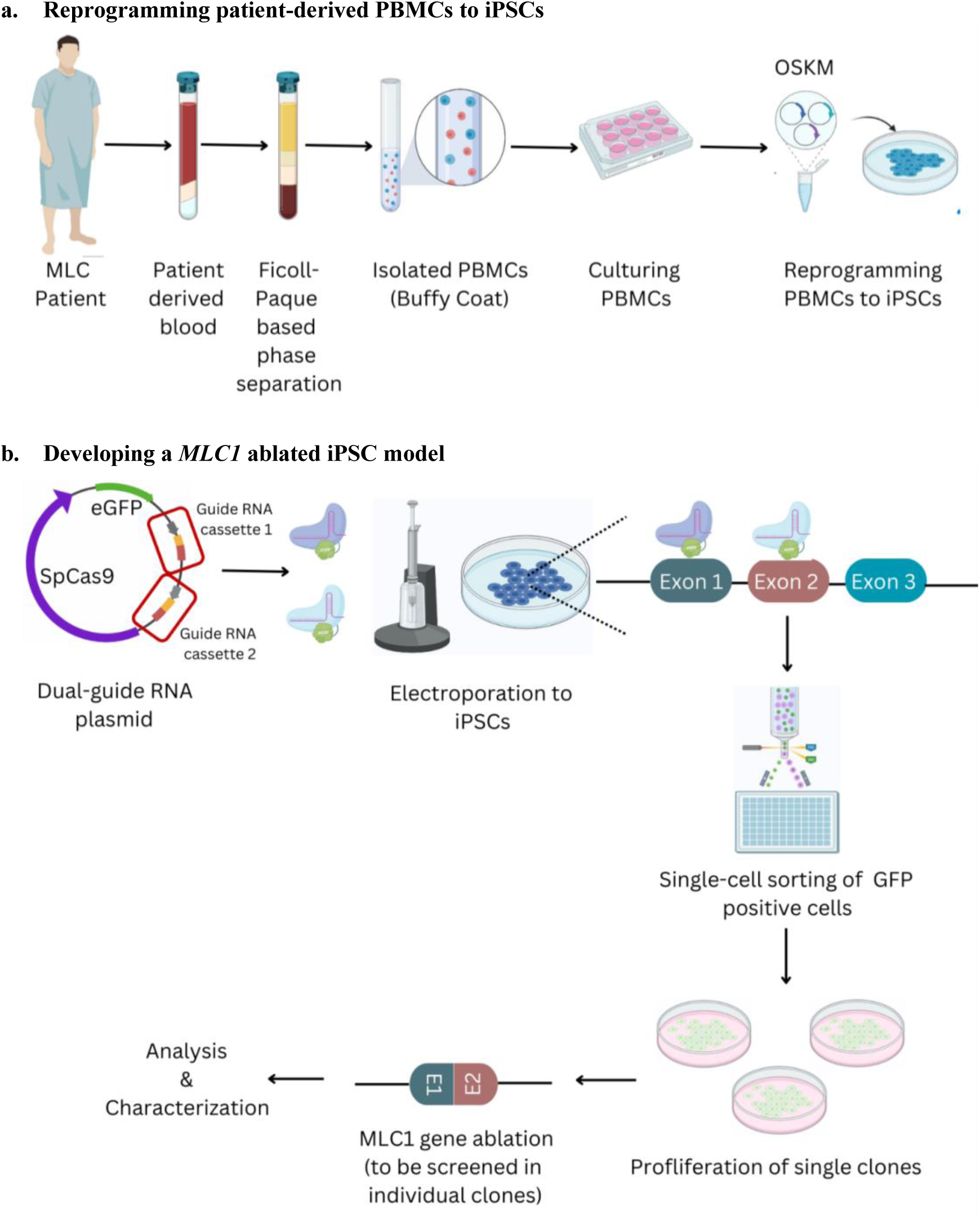
Schematic representation of methodology utilized for MLC iPSC model generation. a. Patient-derived PBMCs were reprogrammed to iPSCs and validated for pluripotency. b. A *MLC1* ablated iPSC line was created via CRISPR-based editing of unaffected iPSCs and clonal isolation and screening.

**Figure 2.**
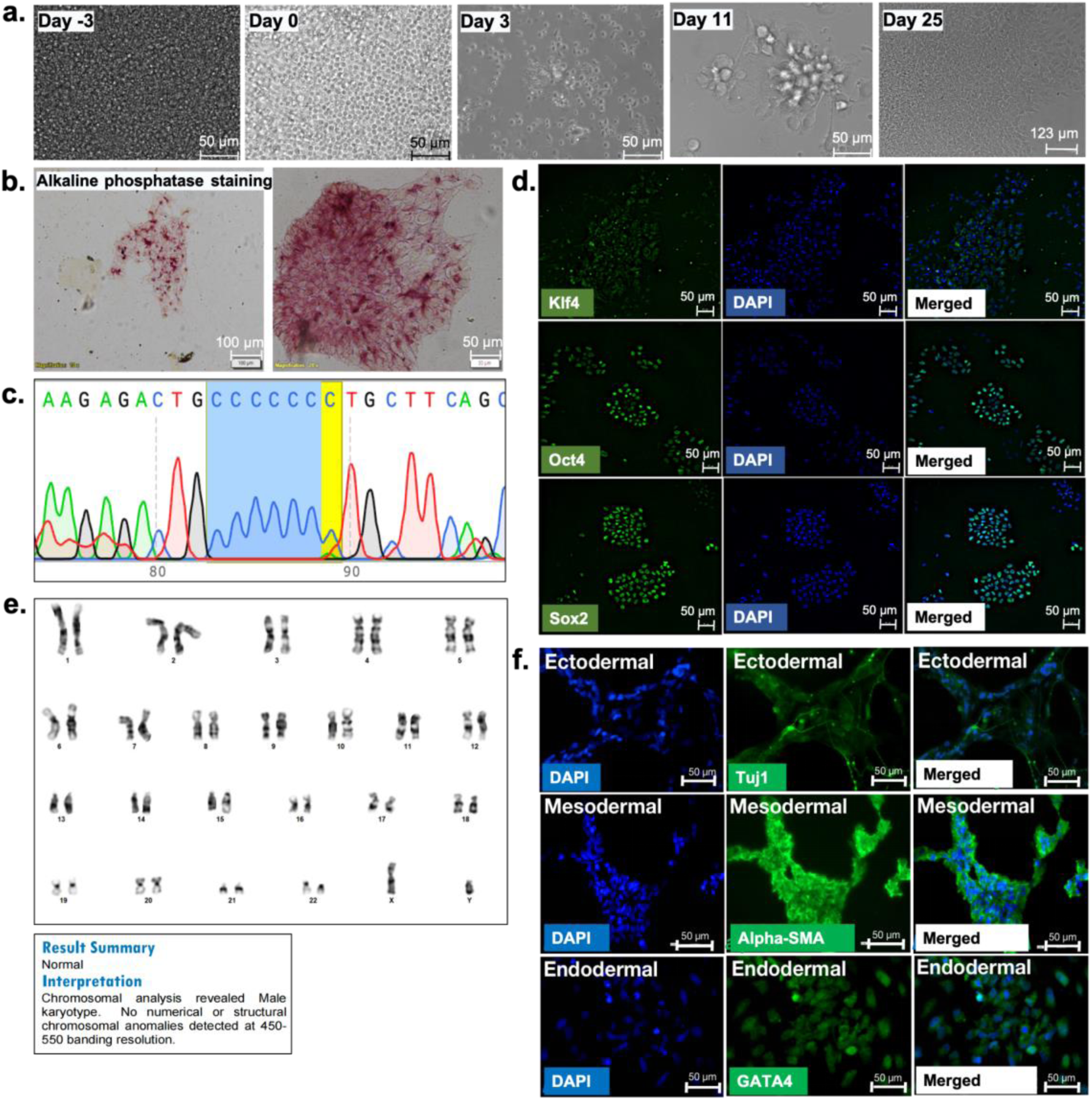
Reprogramming MLC patient PBMCs to iPSCs. a. Sequential progression of patient-derived blood cells reprogrammed to iPSCs. b. Derived-iPSCs were alkaline phosphatase positive. c. Sanger sequencing confirmation of retention of *MLC1* variant (132dupC) in iPSCs. d. Immunolabeling for proteins characteristic of pluripotent & proliferating iPSCs. e. G-banding of patient-derived iPSCs for assessment of karyotypic normalcy. f. Immunolabeling for assessment of iPSC trilineage differentiation efficiency.

### Creation of a genetically ablated MLC1 line

Considering the broad clinical phenotypes of MLC, iPSCs derived solely from patients, though invaluable, may not encapsulate the entirety of the disease pathology at a cellular level. To analyze MLC impacts beyond a specific gene variant, we utilized a CRISPR-Cas9 system in a wild-type iPSC line to create a *MLC1* knockout exhibiting completely inactivated *MLC1* and providing a complementary model system for studying the effects of complete loss of MLC1 function. We cloned a guide RNA (gRNA) sequence targeting *MLC1* into a mammalian expression plasmid. This plasmid construct contained a cassette for the expression of two gRNAs, a *Streptococcus pyogenes* derived Cas9 (SpCas9) gene cassette, and a green fluorescent reporter cassette. The two gRNAs were designed to target distinct regions in *MLC1* to enhance the introduction of multiple double-strand breaks (**Figure 1b**). We transfected HEK293T cells with different combinations of gRNAs in the dual gRNA plasmid to test the genome editing efficiency of the plasmid construct in the pool population (**Supplementary Figure S4**). Following electroporation in iPSCs, we carried out single-cell sorting to isolate individual cells for propagation into distinct clones. Upon sufficient proliferation of these clones, we then screened clones using a genotyping-based PCR to assess *MLC1* deletion in the iPSCs. We successfully identified two clones (signified as MLC1 knockout 2 and MLC1 knockout 3) that demonstrated successful Cas9-mediated *MLC1* deletion (**Figure 3a**). Upon identifying the two clones that displayed the desired genotype, we aimed to further investigate whether *MLC1* deletion occurred in a biallelic manner. We utilized an alternative genotyping strategy in which we designed primer pairs that flanked exons 1 and 2 of *MLC1*. In case of genetic ablation in monoallelic manner, we should get bands at the size of 330 bp and 380 bp with different primer sets, whereas if it were to have happened in a biallelic manner, there should be no band at all (as the genetic ablation would have resulted in the deletion of the *MLC1* region, resulting in an inability of one of the primer set to bind and give rise to an amplicon). As per our observations, both iPSC clones did not show any amplification, thereby suggesting potentially successful biallelic knockout of *MLC1* in these iPSC clones (**Figure 3b**). We subsequently performed Sanger sequencing which corroborated our PCR screening, conclusively confirming that a biallelic knockout of *MLC1* had occurred in our iPSC clones (**Figure 3d**). Following confirmation of *MLC1* knockout in our iPSC clones, we proceeded to carry out characterization for pluripotency in *MLC1* deleted iPSCs and isogenic scrambled controls. To evaluate pluripotency, we utilized an immunofluorescence assay for the expression of pluripotency markers OCT4 and SSEA4. *MLC1* knockout clone retained pluripotent protein expression, suggesting *MLC1* loss does not impact pluripotency (**Figure 3c**). We next examined the trilineage differentiation capacity of the MLC1 knockout iPSC line. To achieve this, we induced undirected differentiation of the iPSCs by permitting cellular aggregation and spontaneous differentiation into embryoid bodies over 21 days **(Figure 3e)**. qPCR analyses validated the trilineage differentiation capacity of *MLC1* deleted iPSCs (**Figure 3f**).

**Figure 3.**
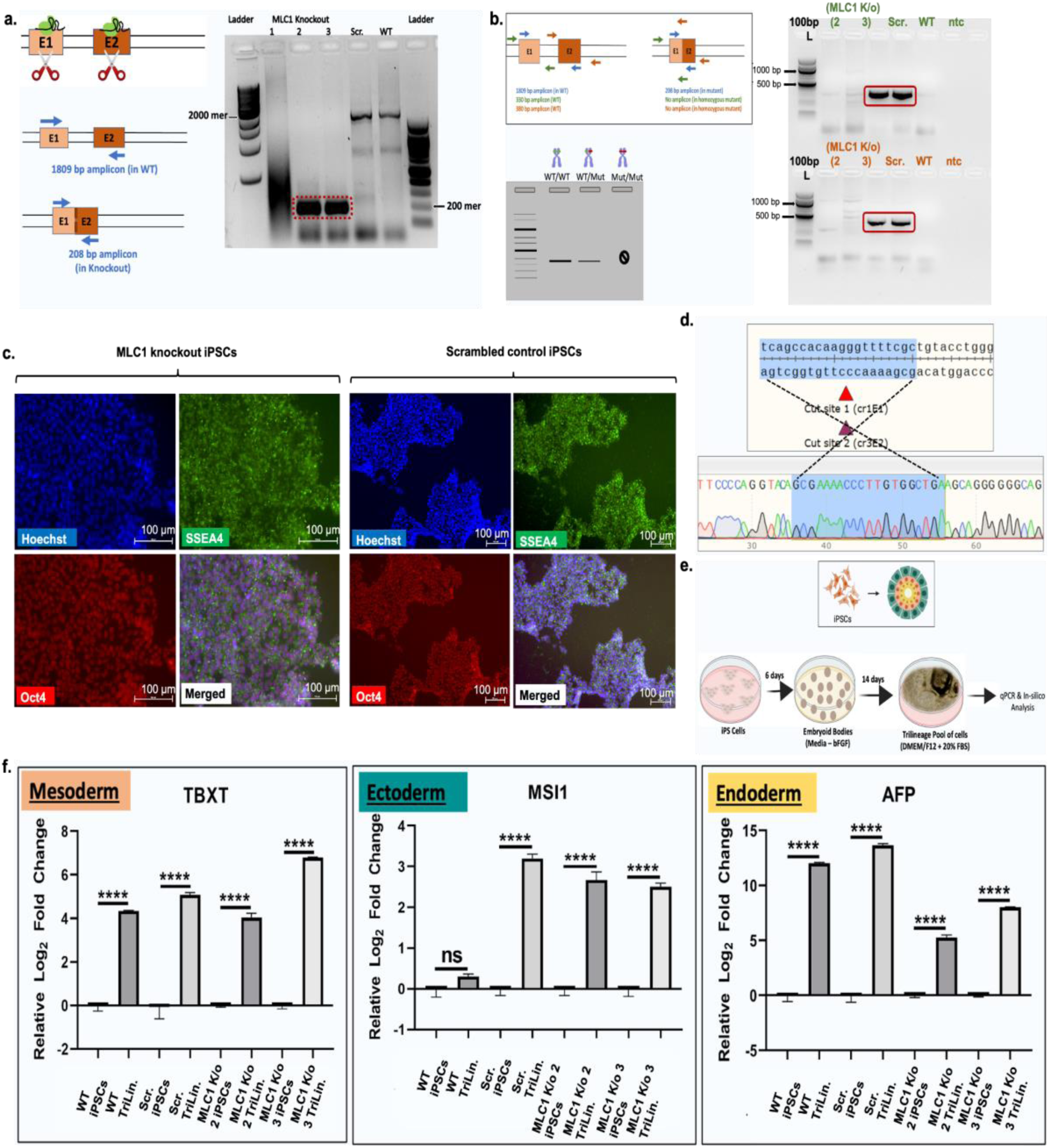
Generation of *MLC1* knockout iPSCs. a. The two ribonucleoprotein complexes from the dual guide plasmid target exon 1 and exon 2 of *MLC1*, resulting in a truncated 200 mer amplicon compared to a ∼2000 mer amplicon in wild-type control. b. PCR based genotyping of clones MLC1 knockout 2 & 3 demonstrate biallelic deletion of *MLC1*. c. *MLC1* ablated iPSCs express markers of pluripotency. d. Representative Sanger sequencing assessment of gRNA targeted *MLC1* in clone *MLC1* knockout 2. e. Schematic of trilineage differentiation of *MLC1* knockout iPSCs. f. *MLC1* knockout iPSCs could be differentiated to all three germ layers as seen by qPCR assay (performed two-tailed, Student’s t-test; **** p-value <0.05)

### Patient-derived and *MLC1* deleted iPSCs successfully differentiate to neural stem cells

To begin to assess the impact of *MLC1* mutation on neurodevelopment, both patient-derived and MLC1 knockout iPSCs were differentiated into neural stem cells (NSCs), a population of self-renewing, multipotent progenitors with the potential to differentiate into neurons, astrocytes, and oligodendrocytes. We speculated that studying *MLC1* knockout NSCs might give us glimpses of potential pathways that eventually lead to disease manifestation in glia. We utilized a detailed differentiation protocol that guides the pluripotent stem cells through sequential stages of differentiation, ultimately leading to the formation of NSCs (**Figure 4a**). The differentiated NSCs that were derived from the patient derived iPSCs (**Figure 4b**), *MLC1* knockout, and scrambled control iPSCs (**Figure 4c**) were immunolabeled with NSC specific antibodies (PAX6, Nestin, SOX1, SOX2). The presence of the respective fluorescence signals in these cell populations provided us with compelling evidence of successful NSC generation.

**Figure 4.**
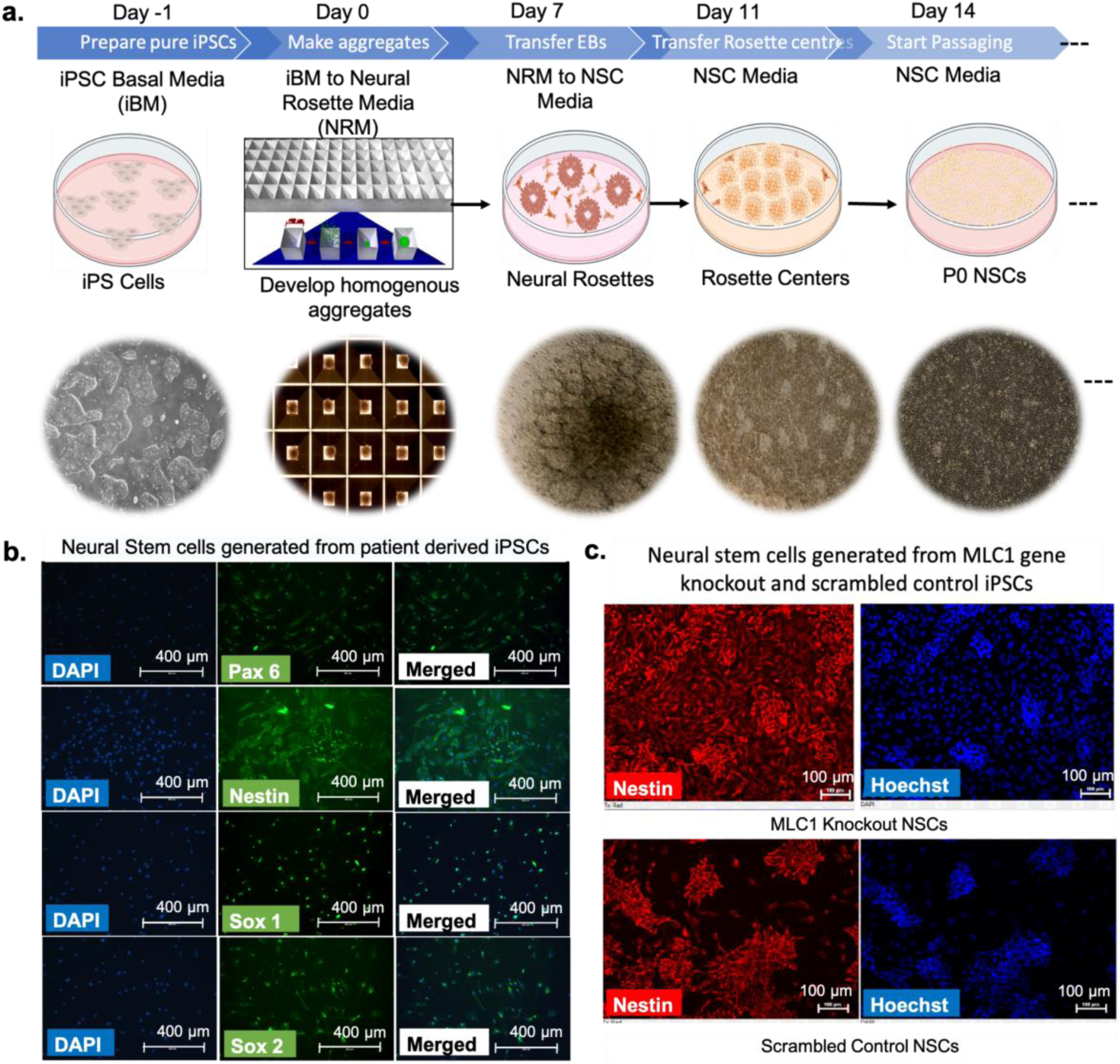
Generation of neural stem cells from MLC patient-derived iPSCs and *MLC1* knockout iPSCs. a. Schematic representation and microscopic (bright-field) depiction of protocol of NSC generation. b. Immunostaining of neural stem cells derived from patient-derived iPSCs. c. Immunostaining of neural stem cells differentiated from *MLC1* and scrambled control iPSCs.

### Transcriptomic profiling of MLC patient-derived and *MLC1* neural stem cells reveals disease-relevant differentially expressed genes

MLC1 expression is predominantly associated with later stages of neural differentiation, specifically when NSCs undergo lineage specification towards astrocytes (Boor et al., 2005)(van der Knaap et al., 2012). However, MLC1 might play an unidentified role in earlier stages of neurodevelopment before astrocyte lineage commitment. NSC transcriptome profiling might offer an opportunity to unveil early pathological markers or identify signaling pathways associated with MLC pathogenesis. Our results revealed a relatively small number of differentially expressed genes (366 total). The differentially expressed genes were primarily associated with neurological disorders including epilepsy (**Figure 5a**; **Supplementary Table S4**). The overall modest number of differentially expressed genes may hint at subtle but crucial alterations in cellular pathways that contribute to the pathogenesis of MLC. The MLC1 gene, however, wasn’t part of this group of differentially expressed genes, potentially indicating the need for an expression above a specific baseline (as seen in astrocytes (**Figure S1**)), to be able to execute a proper disease hallmark. In order to gain more insightful interpretations from our dataset, we classified the differentially expressed genes according to diseases in which they have been implicated (**Supplementary Figure S5**). Surprisingly, the neurological diseases and their corresponding genes identified within our transcriptomic data were all associated with epilepsy, a condition commonly observed in MLC patients. This unexpected link at the NSC stage underscores the potential link between MLC and epileptogenic pathways, suggesting that it might be plausible that MLC’s clinical presentation could be influenced by disturbances in the neuronal circuits associated with epilepsy. Concurrently, we conducted an exhaustive review of experimental, investigational, and approved therapeutics to identify potential drug candidates that could be repurposed for MLC treatment based upon identified epilepsy-associated pathways (**Figure 5b,c**). Beyond identifying signaling pathways of interest to MLC, these data identified within NSCs also suggest MLC pathogenesis may involve both developmental defects and astrocyte dysfunction.

**Figure 5.**
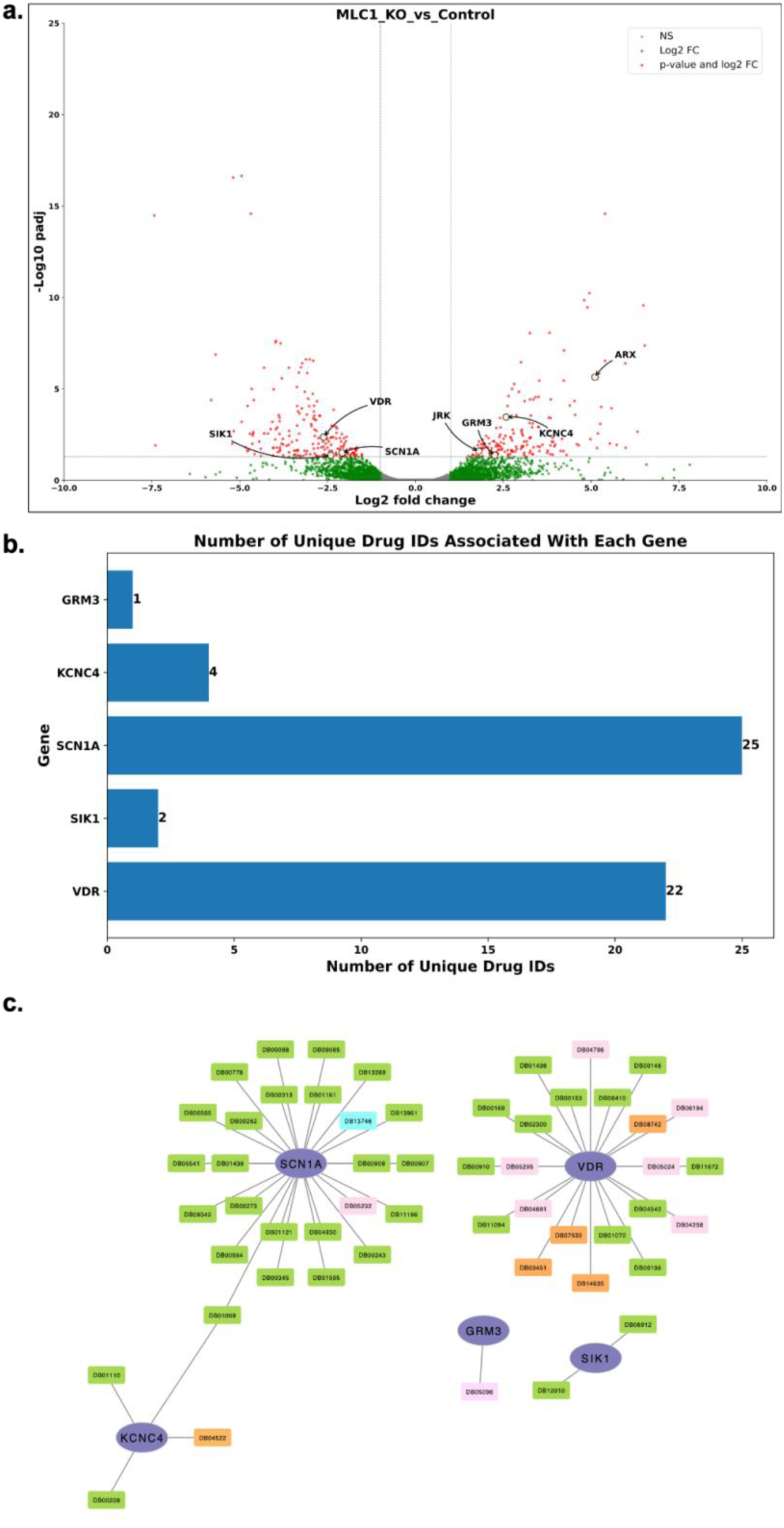
Transcriptomic profiling of neural stem cells derived from *MLC1* deleted iPSCs. a. Volcano plot depicting differentially expressed genes in *MLC1* knockout NSCs compared to isogenic controls. Epilepsy related genes are highlighted. b. Histogram depicting numerical estimate of a comprehensive list of drugs associated with epilepsy related genes. c. Identification of drugs targeting epilepsy-related genes that might be investigated and repurposed for MLC therapy. Green colored drugs are the approved category of drugs, pink is for investigational drugs, blue is for approved+experimental, orange is for experimental drug category, all for the purple colored genes.

### *MLC1* knockout iPSCs successfully differentiate to disease-relevant astrocytes

MLC1 has a specific and high expression in astrocytes (Boor et al., 2005)(van der Knaap et al., 2012) ( **Figure S1**). To model the impact of MLC1 on astrocyte biology, we followed a standardized protocol to differentiate NSCs derived from control, MLC patient-derived NSCs and *MLC1* knockout iPSCs to an astrocyte lineage (Soubannier et al., 2020). This optimization included a series of procedures and conditions tailored to foster the appropriate differentiation of these distinct NSC lines into astrocytes (**Figure 6a**). In the process of astrocyte differentiation, we consistently monitored for morphological changes as a preliminary measure of progression and the success of the differentiation process. By D30, we observed that the cells had begun to express characteristic astrocyte markers, including glial fibrillary acidic protein (GFAP), indicating their successful differentiation into astrocytes. In addition, the MLC1 protein also began expressing in the cells. However, as predicted, the MLC signal was seemingly diminished in the MLC1 gene knockout cell line (**Figure 6b**).

**Figure 6.**
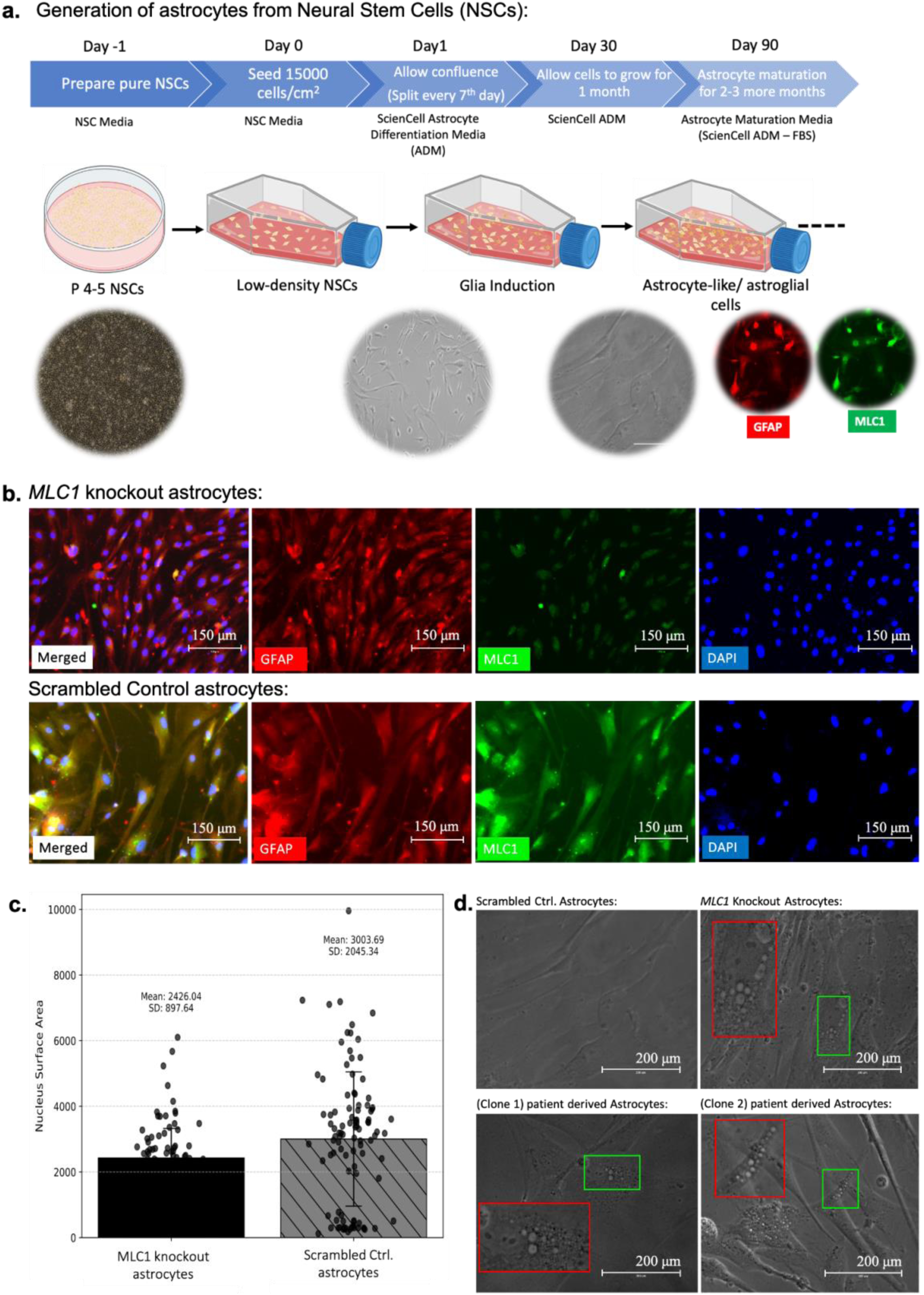
*MLC1* differentiated astrocytes exhibit a vacuolation phenotype characteristic of MLC. a. Schematic representation and microscopic (bright-field) depiction of the protocol used for astrocyte generation. b. MLC1 expression is ablated in *MLC1* knockout astrocytes but highly expressed in controls. c. The cellular nuclei of *MLC1* knockout astrocytes have a smaller surface area compared to scrambled control (Mann-Whitney U test; p-value= 0.0017 ; n=101). d. *MLC1* knockout and patient-derived astrocyte cell lines displayed vacuoles (red inset shows magnification of the area enclosed by green inset), as previously observed in *Mlc1* rat astrocytes (Dubey et al., 2014).

During the course of differentiation, we noted the emergence of vacuolation across the astrocyte cell body of both MLC patient-derived and *MLC1* knockout astrocytes. This phenotypic characteristic, a hallmark of MLC pathology, was distinctly absent in control astrocytes (**Figure 6d**). The appearance of these vacuoles is noteworthy, as it mirrors the phenotype observed in patient brain tissue. Cellular vacuoles can be acidic in nature, such as lysosomes, while others can be neutral or basic. To rule out that morphological MLC vacuoles are lysosomes, we examined if the vacuoles that we observe in MLC astrocytes are acidic. We used LysoTrackerRED, a specific and widely-used fluorescent probe designed for the purpose of labelling and tracking acidic organelles within live cells. Our data revealed that there is little to no colocalization between the LysoTrackerRED dye and the vacuoles within our cells (**Figure S7a,b**). This evidence refutes that a major lot of the observed vacuoles are acidic in nature, thereby ruling out the possibility of them being lysosomal or other acidic vesicles. The recapitulation of this disease-specific vacuolar phenotype strongly affirms that our MLC iPSCs represent a robust model for MLC. Parallely, we observed that the cellular and nuclear size of the *MLC1* knockout iPSCs was shrunken in size as compared to the scrambled control iPSCs (**Figure S6, Figure 6c**).

## DISCUSSION

MLC is a disease that manifests due to mutations in *MLC1* or *GlialCAM* in an autosomal recessive or an autosomal dominant manner. To better delineate MLC pathogenesis, we developed novel iPSC models of MLC due to either a specific *MLC1* mutation (132dupC) found in the Indian Agrawal community or using CRISPR/Cas9 system to create *MLC1* knockout iPSCs. Upon validation of the iPSC lines via multiple parameters to affirm their iPSC status, we also differentiated these cells into disease-relevant neural lineages. We performed transcriptomic profiling of *MLC1* mutant and control neural derivatives, uncovering >350 differentially expressed genes associated with neurological disease including epilepsy. During the course of astrocyte differentiation, we also found vacuolation within the cell bodies of MLC patient-derived and *MLC1* astrocytes, a feature absent from isogenic controls. These data demonstrate that derived MLC iPSCs represent a valid model system capable of recapitulating key hallmarks of MLC disease.

To study MLC pathophysiology, various animal models have been developed. *MLC1* and *GlialCAM* knockouts, including several knockout (KO) and knock-in (KI) models, have been created in mice (Hoegg-Beiler et al., 2014; Dubey et al., 2015; Sugio et al., 2017; Bugiani et al., 2017; Shi et al., 2019; Pérez-Rius et al., 2019), as well as zebrafish (Sirisi et al., 2014; Pérez-Rius et al., 2019). These models share similarities with human MLC patients, such as increased brain water content and intramyelinic vacuoles, detectable by MRI and histological methods (van der Knaap et al., 1996). However, when compared with humans, many major differences exist, including the timing of MRI-detectable defects that occur during the initial stages of life in human patients while reducing and gaining stability at later stages (Teijido et al., 2007; Gilbert et al., 2019). Also, there have been observable regional variations in vacuolization. For human subjects, the subcortical white matter region sustains brain edema. Within MLC mouse models, the pathological defects are mainly observed within the cerebellum (Teijido et al., 2007; López-Hernández et al., 2011a). There are also substantial cellular and functional disparities that exist between human and rodent astrocytes (Oberheim NA et. al., 2009). The human brain exhibits a higher astrocyte-to-neuron ratio compared to rodents, underscoring the importance of deriving astrocytes from a human source using an effective protocol to study human-specific disease states. To circumvent these issues and to complement animal model based studies, there have been studies conducted on lymphoblast cell lines and human monocytes or their derived macrophages from wild-type and MLC patients (Duarri et al., 2008; Petrini et al., 2013). In this scenario, the patient-derived and genetically ablated iPSC based models of MLC disease would add further to the repertoire of MLC disease research and understanding of the disease pathology. The emergence of MLC-related gene expression in early stage neural stem cells and the prevalence of disease-like hallmarks (vacuolation) in differentiated astrocytes provides a glimpse of MLC disease progression during development.

Based upon the signaling pathway changes we have identified, it may be possible to repurpose available investigational or approved drugs targeting epilepsy-related genes for potential therapeutics for MLC. Similar drug screening studies in iPSC-derived neural models have been performed for a variety of rare and common neurological diseases. Lee et al. (2009) modeled the pathogenesis and treatment of familial dysautonomia with derived neural crest precursors from patient-specific iPSCs. Marchetto et al. (2010) modeled neural development and treatment strategies for Rett syndrome as a model of autism spectrum disorder, using patient iPSCs-derived neurons. Yahata et al. (2011) developed an anti-Aβ drug screening platform using iPSC-derived neurons to treat Alzheimer’s disease. Daozhan Y et al., 2014 used patient-derived Niemann–Pick disease type C neural stem cells as a model system for evaluation of drug efficacy and study of the disease pathogenesis. Wang et al. (2017) and Kondo et al. (2017) developed a robust high-content screening assay to identify compounds that reduce tau levels and toxic Aβ levels in iPSC derived neurons, as a potential therapy for Alzheimer’s disease. Similar to these published studies, we propose that our MLC iPSC models and differentiated neural derivatives represent an ideal cellular platform for screening drugs and small molecules which inhibit hallmarks of MLC disease for eventual translation to MLC patients.

In conclusion, we have developed well characterized MLC patient-derived and *MLC1* mutant iPSC models of Megalencephalic Leukoencephalopathy with subcortical cysts. These cellular models can serve as a platform for a variety of screening assays mediated by chemical and genetic interventions, with a strong potential of discovery of a therapeutic solution for MLC and to help better understand the pathogenesis of this disease. In particular, a detailed analysis of the hallmark MLC vacuolation and in-depth characterization of MLC iPSC-derived astrocytes can serve as a phenotypic assay for drug screening platforms or gene augmentation assays to develop or repurpose drugs to improve the lives of MLC patients.

## Supporting information

https://holt-sc.glialab.org/sc/

## LIST OF ABBREVIATIONS

132dupC: MLC1 gene mutation with a cytosine duplication, found in the Agrawal cohort with MLC.
22qtel: Telomeric end of long arm (q) of chromosome 22.
AFP: Alfafeto protein
Alpha-SMA: Smooth muscle alpha-actin
ARX: Aristaless-related homeobox
bFGF: Basic fibroblast growth factor
ClC2: Chloride channel 2 gene
CNS: Central Nervous System
CRISPR: Clustered Regularly Interspaced Short Palindromic Repeats
DMEM: Dulbecco’s Modified Eagle Medium
DNA: Deoxyribonucleic Acid
DSB: Double Strand Breaks
EGF: Epidermal growth factor
en1FnCas9: enhanced variant of FnCas9 protein
ER: Endoplasmic Reticulum
ERAD: Endoplasmic Reticulum Associated Degradation
FACS: Fluorescence Activated Cell Sorting
FLT3: fms-like tyrosine kinase 3
FnCas9: Cas9 variant derived from the bacterial species *Francisella novicida*
G-banding: Giemsa banding
GFAP: Glial Fibrillary Acidic Protein
GFP: Green Fluorescent Protein
GlialCAM: Hepatic and Glial Cell adhesion molecule
gq-PCR: Genomic qPCR
GRM3: Glutamate Metabotropic Receptor 3
gRNA: guide RNA
HDR: Homology Directed Repair
HEK293: Human embryonic Kidney
HEK293T: Human embryonic Kidney 293T
HepaCAM: Hepatic and Glial Cell adhesion molecule
IL-3: Interleukin 3
IL-6: Interleukin 6
indel: insertion/deletion
iPSC: induced Pluripotent Stem Cells
IVC: *In-vitro* cleavage
KCNC4: Potassium Voltage-Gated Channel Subfamily C Member 4
KEGG: Kyoto Encyclopedia of Genes and Genomes
Kir 4.1: Inwardly rectifying potassium
Klf4: Krüppel-like factor 4
KO: Knockout
MLC: Megalencephalic Leukoencephalopathy with subcortical Cysts
MLC1: Megalencephalic Leukoencephalopathy with subcortical Cysts 1
MRI: Magnetic Resonance Imaging
mRNA: messenger RNA
MSI1: Musashi RNA Binding Protein 1
MwoI: MwoI gene from *Methanobacterium wolfeii*
Na K-ATPase: Sodium Potassium Adenosine triphosphate
NGS: Next generation sequencing
NPC: Neural Progenitor Cells
NSC: Neural stem cells
Oct-04: Octamer-binding transcription factor 4
OSKM: Yamanaka factors (Oct3/4, Sox2, Klf4,c-Myc)
PBMC: peripheral blood mononuclear cells
PBS: Phosphate buffer saline
PCR: Polymerase Chain Reaction
PDL: Poly-D-Lysine
PINK1: PTEN induced putative kinase 1
PLO: Poly-L-Ornithine
qPCR: quantitative polymerase chain reaction
RFLP: Restriction Fragment Length Polymorphism
RNA: Ribonucleic Acid
RNP: Ribonucleoprotein
SCF: Stem Cell factor
SCN1A: Sodium channel protein type 1 subunit alpha
SDM: Site-Directed Mutagenesis
sgRNA: single guide RNA
SIK1: Salt Inducible Kinase 1
siRNA: Small interfering RNA
Sox2: SRY-Box Transcription Factor 2
SpCas9: Cas9 variant derived from the bacterial species Streptococcus pyogenes
SSEA4: Stage-specific embryonic antigen-4
TBXT: T-box transcription factor T
TRPV4: Transient receptor potential vanilloid-type 4
V-ATPase: Vacuolar-type ATPase
VDR: Vitamin D receptor
WHO: World Health Organisation
WT: Wild Type
ZO-1: Zonula occludens-1

## ACKNOWLEDGEMENTS

This study was supported by the Council for Scientific and Industrial Research (CSIR), India and the Department of Biotechnology, India. We acknowledge Dr. Sivaprakash Ramalingam’s lab for the training in somatic cell reprogramming, Dr. Sonali Sengupta for assistance with pluripotent stem cell culture practices and neural differentiation assays, and Dr. Manikandan Subhramanian for providing anti-SMA antibody. Some images were drawn with Biorender (Registered under CSIR-IGIB) and MS powerpoint.

## AUTHOR CONTRIBUTIONS

SS conceptualized this study, conducted the experiments, analyzed the data, prepared the figures and wrote the manuscript. All authors assisted in editing of the manuscript and approved of the final manuscript version for publication. HS & RR assisted in maintenance of the cell lines when in culture. HR performed the alkaline phosphatase assay at AIIMS. VB performed the analysis of transcriptomic data and prepared relevant figures. PD helped in analysis of deep sequencing of MLC1 gene amplicon via NGS (not included in this study yet). AR purified the Cas9 protein used in this study. KRF assisted with iPSC culture, directed neural differentiations, and associated practices. DC conceptualized this study, established collaborations and acquired funding, along with MRC, NG, RS, SM, KRF, and MK.

## AVAILABILITY OF DATA AND MATERIALS

All data generated or analysed during this study are included in this article.

## DECLARATIONS

### Ethics approval and consent to participate

The present study was approved by the Institute Ethics Committee (IEC) as well as the Institutional Committee for Stem Cell Research (IC-SCR). Written informed consent was obtained from the MLC patient and blood was collected at AIIMS. This work was sanctioned by DBT under the sanction order no.: BT/25847/GET/119/164/2017.

### Consent for publication

N/A

### Data availability statement

The authors confirm that the data supporting the findings of this study are available within the article [and/or] its supplementary materials. However, some elements of the data (related to patient samples) are not publicly available due to their containing information that could compromise the privacy of research participant(s).

## REFERENCES

Abdel-Salam, G. M. H., Abdel-Hamid, M. S., Ismail, S. I., Hosny, H., Omar, T., Effat, L., Aglan, M. S., Temtamy, S. A., & Zaki, M. S. (2016). Megalencephalic leukoencephalopathy with cysts in twelve Egyptian patients: Novel mutations in MLC1 and HEPACAM and a founder effect. Metabolic Brain Disease, 31(5), 1171–1179. 10.1007/s11011-016-9861-7

Acharya, S., Mishra, A., Paul, D., Ansari, A. H., Azhar, Mohd., Kumar, M., Rauthan, R., Sharma, N., Aich, M., Sinha, D., Sharma, S., Jain, S., Ray, A., Jain, S., Ramalingam, S., Maiti, S., & Chakraborty, D. (2019). Francisella novicida Cas9 interrogates genomic DNA with very high specificity and can be used for mammalian genome editing. Proceedings of the National Academy of Sciences, 116(42), 20959–20968. 10.1073/pnas.1818461116

Ambrosini, E., Serafini, B., Lanciotti, A., Tosini, F., Scialpi, F., Psaila, R., Raggi, C., Di Girolamo, F., Petrucci, T. C., & Aloisi, F. (2008). Biochemical characterization of MLC1 protein in astrocytes and its association with the dystrophin–glycoprotein complex. Molecular and Cellular Neuroscience, 37(3), 480–493. 10.1016/j.mcn.2007.11.003

Batla A, Pandey S, Nehru R. Megalencephalic leukoencephalopathy with subcortical cysts: A report of four cases. J Pediatr Neurosci. 2011 Jan;6(1):74–7. doi: 10.4103/1817-1745.84416

Ben-Zeev, B., Gross, V., Kushnir, T., Shalev, R., Hoffman, C., Shinar, Y., Pras, E., & Brand, N. (2001). Vacuolating megalencephalic leukoencephalopathy in 12 Israeli patients. Journal Of Child Neurology, 16(02), 093. 10.2310/7010.2001.6983

Brennand KJ, Simone A, Jou J, Gelboin-Burkhart C, Tran N, Sangar S, Li Y, Mu Y, Chen G, Yu D, McCarthy S, Sebat J, Gage FH. Modelling schizophrenia using human induced pluripotent stem cells. Nature. 2011 May 12;473(7346):221-5. doi: 10.1038/nature09915

Brignone, M. S., Lanciotti, A., Macioce, P., Macchia, G., Gaetani, M., Aloisi, F., Petrucci, T. C., & Ambrosini, E. (2010). The β1 subunit of the Na,K-ATPase pump interacts with megalencephalic leucoencephalopathy with subcortical cysts protein 1 (MLC1) in brain astrocytes: New insights into MLC pathogenesis. Human Molecular Genetics, 20(1), 90–103. 10.1093/hmg/ddq435

Bugiani, M., Dubey, M., Breur, M., Postma, N. L., Dekker, M. P., Ter Braak, T., et al. (2017). Megalencephalic leukoencephalopathy with cysts: the Glialcam-null mouse model. Ann. Clin. Transl. Neurol. 4, 450–465. doi: 10.1002/acn3.405

Capdevila-Nortes, X., López-Hernández, T., Apaja, P. M., López de Heredia, M., Sirisi, S., Callejo, G., Arnedo, T., Nunes, V., Lukacs, G. L., Gasull, X., & Estévez, R. (2013). Insights into MLC pathogenesis: GlialCAM is an MLC1 chaperone required for proper activation of volume-regulated anion currents. Human Molecular Genetics, 22(21), 4405–4416. 10.1093/hmg/ddt290

Chen, J. S., Dagdas, Y. S., Kleinstiver, B. P., Welch, M. M., Sousa, A. A., Harrington, L. B., Sternberg, S. H., Joung, J. K., Yildiz, A., & Doudna, J. A. (2017). Enhanced proofreading governs CRISPR–Cas9 targeting accuracy. Nature, 550(7676), 407–410. 10.1038/nature24268

Chen, Y., & Swanson, R. A. (2003). Astrocytes and brain injury. Journal of Cerebral Blood Flow & Metabolism, 23(2), 137–149. 10.1097/01.wcb.0000044631.80210.3c

Chen, Y., Zheng, Y., Kang, Y., Yang, W., Niu, Y., Guo, X., Tu, Z., Si, C., Wang, H., Xing, R., Pu, X., Yang, S.- H., Li, S., Ji, W., & Li, X.-J. (2015). Functional disruption of the dystrophin gene in rhesus monkey using CRISPR/Cas9. Human Molecular Genetics, 24(13), 3764–3774. 10.1093/hmg/ddv120

Chertow, D. S. (2018). Next-generation diagnostics with CRISPR. Science, 360(6387), 381–382. 10.1126/science.aat4982

Choi, S. A., Kim, S. Y., Yoon, J., Choi, J., Park, S. S., Seong, M.-W., Kim, H., Hwang, H., Choi, J. E., Chae, J. H., Kim, K. J., Kim, S., Lee, Y.-J., Nam, S. O., & Lim, B. C. (2017). A Unique Mutational Spectrum of MLC1 in Korean Patients With Megalencephalic Leukoencephalopathy With Subcortical Cysts: P.Ala275Asp Founder Mutation and Maternal Uniparental Disomy of Chromosome 22. Annals of Laboratory Medicine, 37(6), 516–521. 10.3343/alm.2017.37.6.516

Cong, L., Ran, F. A., Cox, D., Lin, S., Barretto, R., Habib, N., Hsu, P. D., Wu, X., Jiang, W., Marraffini, L. A., & Zhang, F. (2013).Multiplex genome engineering using crispr/cas systems. Science, 339(6121), 819–823. 10.1126/science.1231143

Damle, Y. B. (1981). India: V. C. Channa: Caste: Identity and Continuity. B. R. Publishing Co., New Delhi, 1979, x, 180p., Rs. 45. India Quarterly: A Journal of International Affairs, 37(1), 129–129. 10.1177/097492848103700129

Duarri, A., Lopez de Heredia, M., Capdevila-Nortes, X., Ridder, M. C., Montolio, M., López-Hernández, T., Boor, I., Lien, C.-F., Hagemann, T., Messing, A., Gorecki, D. C., Scheper, G. C., Martínez, A., Nunes, V., van der Knaap, M. S., & Estévez, R. (2011). Knockdown of MLC1 in primary astrocytes causes cell vacuolation: A MLC disease cell model. Neurobiology of Disease, 43(1), 228–238. 10.1016/j.nbd.2011.03.015

Duarri, A., Teijido, O., Lopez-Hernandez, T., Scheper, G. C., Barriere, H., Boor, I., Aguado, F., Zorzano, A., Palacin, M., Martinez, A., Lukacs, G. L., van der Knaap, M. S., Nunes, V., & Estevez, R. (2008). Molecular pathogenesis of megalencephalic leukoencephalopathy with subcortical cysts: Mutations in MLC1 cause folding defects. Human Molecular Genetics, 17(23), 3728–3739. 10.1093/hmg/ddn269

Dubey, M., Bugiani, M., Ridder, M. C., Postma, N. L., Brouwers, E., Polder, E., Jacobs, J. G., Baayen, J. C., Klooster, J., Kamermans, M., Aardse, R., de Kock, C. P. J., Dekker, M. P., van Weering, J. R. T., Heine, V. M., Abbink, T. E. M., Scheper, G. C., Boor, I., Lodder, J. C., …van der Knaap, M. S. (2014). Mice with megalencephalic leukoencephalopathy with cysts: A developmental angle. Annals of Neurology, 77(1), 114–131. 10.1002/ana.24307

Dubey, M., Bugiani, M., Ridder, M. C., Postma, N. L., Brouwers, E., Polder, E., et al. (2015). Mice with megalencephalic leukoencephalopathy with cysts: a developmental angle. Ann. Neurol. 77, 114–131. doi: 10.1002/ana.24307

Gammie, T., Lu, C. Y., & Babar, Z. U.-D. (2015). Access to orphan drugs: A comprehensive review of legislations, regulations and policies in 35 countries. PLOS ONE, 10(10), e0140002. 10.1371/journal.pone.0140002

Gasiunas, G., Barrangou, R., Horvath, P., & Siksnys, V. (2012). Cas9–crRNA ribonucleoprotein complex mediates specific DNA cleavage for adaptive immunity in bacteria. Proceedings of the National Academy of Sciences, 109(39). 10.1073/pnas.1208507109

Gilbert, A., Vidal, X. E., Estévez, R., Cohen-Salmon, M., and Boulay, A.-C. (2019). Postnatal development of the astrocyte perivascular MLC1/GlialCAM complex defines a temporal window for the gliovascular unit maturation. Brain Struct. Funct. 224, 1267–1278. doi: 10.1007/s00429-019-01832-w

Gupta, V., Khadgawat, R., Ng, H. K. T., Kumar, S., Rao, V. R., & Sachdeva, M. P. (2010). Population structure of aggarwals of North India as revealed by molecular markers. Genetic Testing and Molecular Biomarkers, 14(6), 781–785. 10.1089/gtmb.2010.0095

Gorospe, J. R., Singhal, B. S., Kainu, T., Wu, F., Stephan, D., Trent, J., Hoffman, E. P., & Naidu, S. (2004). Indian Agarwal megalencephalic leukodystrophy with cysts is caused by a commonMLC1mutation. Neurology, 62(6), 878–882. 10.1212/01.wnl.0000115106.88813.5b

Hirano, H., Gootenberg, J. S., Horii, T., Abudayyeh, O. O., Kimura, M., Hsu, P. D., Nakane, T., Ishitani, R., Hatada, I., Zhang, F., Nishimasu, H., & Nureki, O. (2016). Structure and Engineering of Francisella novicida Cas9. Cell, 164(5), 950–961. 10.1016/j.cell.2016.01.039

Hoegg-Beiler, M. B., Sirisi, S., Orozco, I. J., Ferrer, I., Hohensee, S., Auberson, M., Gödde, K., Vilches, C., de Heredia, M. L., Nunes, V., Estévez, R., & Jentsch, T. J. (2014). Disrupting MLC1 and GlialCAM and ClC-2 interactions in leukodystrophy entails glial chloride channel dysfunction. Nature Communications, 5(1). 10.1038/ncomms4475

Ilja Boor, P. K., Groot, K. de, Mejaski-Bosnjak, V., Brenner, C., van der Knaap, M. S., Scheper, G. C., & Pronk, J. C. (2006). Megalencephalic leukoencephalopathy with subcortical cysts: An update and extended mutation analysis of MLC1. Human Mutation, 27(6), 505–512. 10.1002/humu.20332

Jinek, M., Chylinski, K., Fonfara, I., Hauer, M., Doudna, J. A., & Charpentier, E. (2012). A programmable dual-rna–guided DNA endonuclease in adaptive bacterial immunity. Science, 337(6096), 816–821. 10.1126/science.1225829

Kater, M. S. J., Baumgart, K. F., Badia-Soteras, A., Heistek, T. S., Carney, K. E., Timmerman, A. J., van Weering, J. R. T., Smit, A. B., van der Knaap, M. S., Mansvelder, H. D., Verheijen, M. H. G., & Min, R. (2023). A novel role for MLC1 in regulating astrocyte–synapse interactions. Glia, 71(7), 1770–1785. 10.1002/glia.24368

Kim, J. B., Zaehres, H., Wu, G., Gentile, L., Ko, K., Sebastiano, V., Araúzo-Bravo, M. J., Ruau, D., Han, D. W., Zenke, M., & Schöler, H. R. (2008). Pluripotent stem cells induced from adult neural stem cells by reprogramming with two factors. Nature, 454(7204), 646–650. 10.1038/nature07061

Knaap, M. S. van der, Abbink, T. E., & Min, R. (2018, March 29). Megalencephalic leukoencephalopathy with subcortical cysts. NCBI Bookshelf. https://www.ncbi.nlm.nih.gov/books/NBK1535/

Lanciotti, A., Brignone, M. S., Molinari, P., Visentin, S., De Nuccio, C., Macchia, G., Aiello, C., Bertini, E., Aloisi, F., Petrucci, T. C., & Ambrosini, E. (2012). Megalencephalic leukoencephalopathy with subcortical cysts protein 1 functionally cooperates with the TRPV4 cation channel to activate the response of astrocytes to osmotic stress: Dysregulation by pathological mutations. Human Molecular Genetics, 21(10), 2166–2180. 10.1093/hmg/dds032

Lee G., Ramirez C.N., Kim H., Zeltner N., Liu B., Radu C., Bhinder B., Kim Y.J., Choi I.Y., Mukherjee-Clavin B.. et al. (2012) Large-scale screening using familial dysautonomia induced pluripotent stem cells identifies compounds that rescue IKBKAP expression. Nature Biotechnol., 30, 1244–1248. doi: 10.1038/nbt.2435

Leegwater, P. A. J., Yuan, B. Q., van der Steen, J., Mulders, J., Könst, A. A. M., Boor, P. K. I., Mejaski-Bosnjak, V., van der Maarel, S. M., Frants, R. R., Oudejans, C. B. M., Schutgens, R. B. H., Pronk, J. C., & van der Knaap, M. S. (2001). Mutations of MLC1 (KIAA0027), encoding a putative membrane protein, cause megalencephalic leukoencephalopathy with subcortical cysts. The American Journal of Human Genetics, 68(4), 831–838. 10.1086/319519

Liu, H., & Zhang, S.-C. (2011). Specification of neuronal and glial subtypes from human pluripotent stem cells. Cellular and Molecular Life Sciences, 68(24), 3995–4008. 10.1007/s00018-011-0770-y

Liu, J., Gao, C., Chen, W., Ma, W., Li, X., Shi, Y., Zhang, H., Zhang, L., Long, Y., Xu, H., Guo, X., Deng, S., Yan, X., Yu, D., Pan, G., Chen, Y., Lai, L., Liao, W., & Li, Z. (2016). CRISPR/Cas9 facilitates investigation of neural circuit disease using human iPSCs: Mechanism of epilepsy caused by an SCN1A loss-of-function mutation. Translational Psychiatry, 6(1), e703–e703. 10.1038/tp.2015.203

Liu X, Li C, Zheng K, Zhao X, Xu X, Yang A, Yi M, Tao H, Xie B, Qiu M, Yang J. Chromosomal aberration arises during somatic reprogramming to pluripotent stem cells. Cell Div. 2020 Nov 3;15(1):12. doi: 10.1186/s13008-020-00068-z

López-Hernández, T., Ridder, M. C., Montolio, M., Capdevila-Nortes, X., Polder, E., Sirisi, S., Duarri, A., Schulte, U., Fakler, B., Nunes, V., Scheper, G. C., Martínez, A., Estévez, R., & van der Knaap, M. S. (2011). Mutant glialcam causes megalencephalic leukoencephalopathy with subcortical cysts, benign familial macrocephaly, and macrocephaly with retardation and autism. The American Journal of Human Genetics, 88(4), 422–432. 10.1016/j.ajhg.2011.02.009

López-Hernández, T., Sirisi, S., Capdevila-Nortes, X., Montolio, M., Fernández-Dueñas, V., Scheper, G. C., van der Knaap, M. S., Casquero, P., Ciruela, F., Ferrer, I., Nunes, V., & Estévez, R. (2011). Molecular mechanisms of MLC1 and GLIALCAM mutations in megalencephalic leukoencephalopathy with subcortical cysts. Human Molecular Genetics, 20(16), 3266–3277. 10.1093/hmg/ddr238

Mali, P., Yang, L., Esvelt, K. M., Aach, J., Guell, M., DiCarlo, J. E., Norville, J. E., & Church, G. M. (2013). RNA-Guided human genome engineering via cas9. Science, 339(6121), 823–826. 10.1126/science.1232033

Marchetto M.C., Carromeu C., Acab A., Yu D., Yeo G.W., Mu Y., Chen G., Gage F.H., Muotri A.R. (2010) A model for neural development and treatment of Rett syndrome using human induced pluripotent stem cells. Cell, 143, 527–539. doi: 10.1016/j.cell.2010.10.016

Masaki, K., Suzuki, S. O., Matsushita, T., Yonekawa, T., Matsuoka, T., Isobe, N., Motomura, K., Wu, X.-M., Tabira, T., Iwaki, T., & Kira, J. (2012). Extensive loss of connexins in Baló’s disease: Evidence for an auto-antibody-independent astrocytopathy via impaired astrocyte–oligodendrocyte/myelin interaction. Acta Neuropathologica, 123(6), 887–900. 10.1007/s00401-012-0972-x

McKeithan W.L., Feyen D.A.M., Bruyneel A.A.N., Okolotowicz K.J., Ryan D.A., Sampson K.J., Potet F., Savchenko A., Gómez-Galeno J., Vu M., et al. Reengineering an Antiarrhythmic Drug Using Patient hiPSC Cardiomyocytes to Improve Therapeutic Potential and Reduce Toxicity. Cell Stem Cell. 2020;27:813–821.e816. doi: 10.1016/j.stem.2020.08.003

Megalencephalic leukoencephalopathy with subcortical cysts - About the Disease. (n.d.). Genetic and Rare Diseases Information Center. Retrieved June 26, 2023, from https://rarediseases.info.nih.gov/diseases/3445/megalencephalic-leukoencephalopathy-with-subcortical-cysts

Ministry of health and family welfare. (n.d.). National policy for treatment of rare diseases. Mohfd.Gov.In. https://main.mohfw.gov.in/sites/default/files/Rare%20Diseases%20Policy%20FINAL.pdf

Montagna, G., Teijido, O., Eymard-Pierre, E., Muraki, K., Cohen, B., Loizzo, A., Grosso, P., Tedeschi, G., Palacín, M., Boespflug-Tanguy, O., Bertini, E., Santorelli, F. M., & Estévez, R. (2006). Vacuolating megalencephalic leukoencephalopathy with subcortical cysts: Functional studies of novel variants inMLC1. Human Mutation, 27(3), 292–292. 10.1002/humu.9407

Mojica, F. J. M., D□ez-Villase□or, C., Garcia-Martinez, J., & Soria, E. (2005). Intervening sequences of regularly spaced prokaryotic repeats derive from foreign genetic elements. Journal of Molecular Evolution, 60(2), 174–182. 10.1007/s00239-004-0046-3

Oberheim NA, Takano T, Han X, He W, Lin JH, Wang F, Xu Q, Wyatt JD, Pilcher W, Ojemann JG, Ransom BR, Goldman SA, Nedergaard M. Uniquely hominid features of adult human astrocytes. J Neurosci. 2009 Mar 11;29(10):3276–87. doi: 10.1523/JNEUROSCI.4707-08.2009

Office, P. (n.d.). Council Recommendation of 8 June 2009 on an action in the field of rare diseases.

Okita, K., Nakagawa, M., Hyenjong, H., Ichisaka, T., & Yamanaka, S. (2008). Generation of mouse induced pluripotent stem cells without viral vectors. Science, 322(5903), 949–953. 10.1126/science.1164270

Orphanet: Diseases list. (n.d.). Retrieved August 3, 2023, from https://www.orpha.net/consor/cgi-bin/Disease_Search_List.php?lng=EN&TAG=A

Pérez-Rius, C., Folgueira, M., Elorza-Vidal, X., Alia, A., Hoegg-Beiler, M. B., Eeza, M. N.H., et al. (2019). Comparison of zebrafish and mice knockouts for megalencephalic leukoencephalopathy proteins indicates that GlialCAM/MLC1 forms a functional unit. Orphanet J. Rare Dis. 14:268. DOI: 10.1186/s13023-019-1248-5

Petrini, S., Minnone, G., Coccetti, M., Frank, C., Aiello, C., Cutarelli, A., Ambrosini, E., Lanciotti, A., Brignone, M. S., D’Oria, V., Strippoli, R., De Benedetti, F., Bertini, E., & Bracci-Laudiero, L. (2013). Monocytes and macrophages as biomarkers for the diagnosis of megalencephalic leukoencephalopathy with subcortical cysts. Molecular and Cellular Neuroscience, 56, 307–321. 10.1016/j.mcn.2013.07.001

Pfrieger, F. W. (2009). Roles of glial cells in synapse development. Cellular and Molecular Life Sciences, 66(13), 2037–2047. 10.1007/s00018-009-0005-7

Rare diseases in the age of health 2.0. (2014). Springer Berlin Heidelberg. 10.1007/978-3-642-38643-5

Ridder, M. C., Boor, I., Lodder, J. C., Postma, N. L., Capdevila-Nortes, X., Duarri, A., Brussaard, A. B., Estévez, R., Scheper, G. C., Mansvelder, H. D., & van der Knaap, M. S. (2011). Megalencephalic leucoencephalopathy with cysts: Defect in chloride currents and cell volume regulation. Brain, 134(11), 3342–3354. 10.1093/brain/awr255

Rudge, J. S. (1993). Astrocyte-Derived neurotrophic factors. In Astrocytes (pp. 267–305). Elsevier. 10.1016/b978-0-12-511370-0.50016-3

Saijo, H., Nakayama, H., Ezoe, T., Araki, K., Sone, S., Hamaguchi, H., Suzuki, H., Shiroma, N., Kanazawa, N., Tsujino, S., Hirayama, Y., & Arima, M. (2003). A case of megalencephalic leukoencephalopathy with subcortical cysts (van der Knaap disease): Molecular genetic study. Brain and Development, 25(5), 362–366. 10.1016/s0387-7604(03)00006-8

Shi, Z., Yan, H.-F., Cao, B.-B., Guo, M.-M., Xie, H., Gao, K., et al. (2019). Identification in Chinese patients with GLIALCAM mutations of megalencephalic leukoencephalopathy with subcortical cysts and brain pathological study on Glialcam knock-in mouse models. World J. Pediatr. 15, 454–464. doi: 10.1007/s12519-019-00284-w

Shao, Y., & McCarthy, K. D. (1994). Plasticity of astrocytes. Glia, 11(2), 147–155. 10.1002/glia.440110209

Singhal, B. S. (2005). Leukodystrophies: Indian scenario. The Indian Journal of Pediatrics, 72(4), 315–318. 10.1007/bf02724013

Singhal, B. S., Gorospe, J. R., & Naidu, S. (2003). Megalencephalic leukoencephalopathy with subcortical cysts. Journal of Child Neurology, 18(9), 646–652. 10.1177/08830738030180091201

Singhal, B. S., Gursahani, R. D., Udani, V. P., & Biniwale, A. A. (1996). Megalencephalic leukodystrophy in an Asian Indian ethnic group. Pediatric Neurology, 14(4), 291–296. 10.1016/0887-8994(96)00048-3

Sirisi, S., Folgueira, M., López-Hernández, T., Minieri, L., Pérez-Rius, C., Gaitán-Peñas, H., Zang, J., Martínez, A., Capdevila-Nortes, X., De La Villa, P., Roy, U., Alia, A., Neuhauss, S., Ferroni, S., Nunes, V., Estévez, R., & Barrallo-Gimeno, A. (2014). Megalencephalic leukoencephalopathy with subcortical cysts protein 1 regulates glial surface localization of GLIALCAM from fish to humans. Human Molecular Genetics, 23(19), 5069–5086. 10.1093/hmg/ddu231

Sohn, Y.-D., Han, J. W., & Yoon, Y. (2012). Generation of induced pluripotent stem cells from somatic cells. In Progress in Molecular Biology and Translational Science (pp. 1–26). Elsevier. 10.1016/b978-0-12-398459-3.00001-0

Steinke, V., Meyer, J., Syagailo, Y. V., Ortega, G., Hameister, H., M□ssner, R., Schmitt, A., & Lesch, K.-P. (2003). The genomic organization of the murine Mlc1 ( Wkl1 , KIAA0027) gene. Journal of Neural Transmission, 110(4), 333–343. 10.1007/s00702-002-0788-2

Sugio, S., Tohyama, K., Oku, S., Fujiyoshi, K., Yoshimura, T., Hikishima, K., et al. (2017). Astrocyte-mediated infantile-onset leukoencephalopathy mouse model. Glia 65, 150–168. doi: 10.1002/glia.23084

Takahashi, K., & Yamanaka, S. (2006). Induction of pluripotent stem cells from mouse embryonic and adult fibroblast cultures by defined factors. Cell, 126(4), 663–676. 10.1016/j.cell.2006.07.024

Teijido, O., Martínez, A., Pusch, M., Zorzano, A., Soriano, E., del Río, J. A., Palacín, M., & Estévez, R. (2004). Localization and functional analyses of the MLC1 protein involved in megalencephalic leukoencephalopathy with subcortical cysts. Human Molecular Genetics, 13(21), 2581–2594. 10.1093/hmg/ddh291

Teijido, O., Casaroli-Marano, R., Kharkovets, T., Aguado, F., Zorzano, A., Palacin, M., et al. (2007). Expression patterns of MLC1 protein in the central and peripheral nervous systems. Neurobiol. Dis. 26, 532–545. doi: 10.1016/j.nbd.2007.01.016

Topçu, M., Gartioux, C., Ribierre, F., Yalçinkaya, C., Tokus, E., Öztekin, N., Beckmann, J. S., Ozguc, M., & Seboun, E. (2000). Vacuoliting megalencephalic leukoencephalopathy with subcortical cysts, mapped to chromosome 22qtel. The American Journal of Human Genetics, 66(2), 733–739. 10.1086/302758

Torres-Ruiz, R., & Rodriguez-Perales, S. (2016). CRISPR-Cas9 technology: Applications and human disease modelling. Briefings in Functional Genomics, 16(1), 4–12. 10.1093/bfgp/elw025

Vadodaria, K. C., Amatya, D. N., Marchetto, M. C., & Gage, F. H. (2018). Modeling psychiatric disorders using patient stem cell-derived neurons: A way forward. Genome Medicine, 10(1). 10.1186/s13073-017-0512-3

Valk, J., Gorospe, J. R., Singhal, B. S., Hoffman, E. P., & Naidu, S. (2004). Indian Agarwal megalencephalic leukodystrophy with cysts is caused by a common MLC1 mutation. Neurology, 63(11), 2197–2197. 10.1212/wnl.63.11.2197-a

van der Knaap, M. S., Barth, P. G., Stroink, H., van Nieuwenhuizen, O., Arts, W. F. M., Hoogenraad, F., & Valk, J. (1995). Leukoencephalopathy with swelling and a discrepantly mild clinical course in eight children. Annals of Neurology, 37(3), 324–334. 10.1002/ana.410370308

van der Knaap, M. S., Boor, I., & Estévez, R. (2012). Megalencephalic leukoencephalopathy with subcortical cysts: Chronic white matter oedema due to a defect in brain ion and water homoeostasis. The Lancet Neurology, 11(11), 973–985. 10.1016/s1474-4422(12)70192-8

van der Laan, L. J. W., De Groot, C. J. A., Elices, M. J., & Dijkstra, C. D. (1997). Extracellular matrix proteins expressed by human adult astrocytes in vivo and in-vitro: An astrocyte surface protein containing the CS1 domain contributes to binding of lymphoblasts. Journal of Neuroscience Research, 50(4), 539–548. 10.1002/(sici)1097-4547(19971115)50:4<539::aid-jnr5>3.0.co;2-f

Wang, X., Cao, C., Huang, J., Yao, J., Hai, T., Zheng, Q., Wang, X., Zhang, H., Qin, G., Cheng, J., Wang, Y., Yuan, Z., Zhou, Q., Wang, H., & Zhao, J. (2016). One-step generation of triple gene-targeted pigs using CRISPR/Cas9 system. Scientific Reports, 6(1). 10.1038/srep20620

Wang C, et al. Scalable production of iPSC-derived human neurons to identify tau-lowering compounds by high-content screening. Stem Cell Rep. 2017;9(4):1221–33.DOI: 10.1016/j.stemcr.2017.08.019

Kondo T, et al. iPSC-based compound screening and in vitro trials identify a synergistic anti-amyloid β combination for Alzheimer’s disease. Cell Rep. 2017;21(8):2304–12. DOI: 10.1016/j.celrep.2017.10.109

Yahata N, Asai M, Kitaoka S, Takahashi K, Asaka I, Hioki H, Kaneko T, Maruyama K, Saido TC, Nakahata T, Asada T, Yamanaka S, Iwata N, Inoue H. Anti-Aβ drug screening platform using human iPS cell-derived neurons for the treatment of Alzheimer’s disease. PLoS One. 2011;6(9):e25788. doi: 10.1371/journal.pone.0025788

Yamanaka, S. (2012). Induced pluripotent stem cells: Past, present, and future. Cell Stem Cell, 10(6), 678–684. 10.1016/j.stem.2012.05.005

Yiş, U., Scheper, G. C., Uran, N., Ünalp, A., Çakmakçı, H., Kurul, S. H., Dirik, E., & van der Knaap, M. S. (2009). P279 Two cases with megalencephalic leukoencephalopathy with subcortical cysts and MLC1 mutations in Turkish population. European Journal of Paediatric Neurology, 13, S108. 10.1016/s1090-3798(09)70337-x

