## Supplementary material for "*MLC1* alteration in iPSCs give rise to disease-like cellular vacuolation phenotype in the astrocyte lineage": https://holt-sc.glialab.org/sc/

### SUPPLEMENTARY INFORMATION

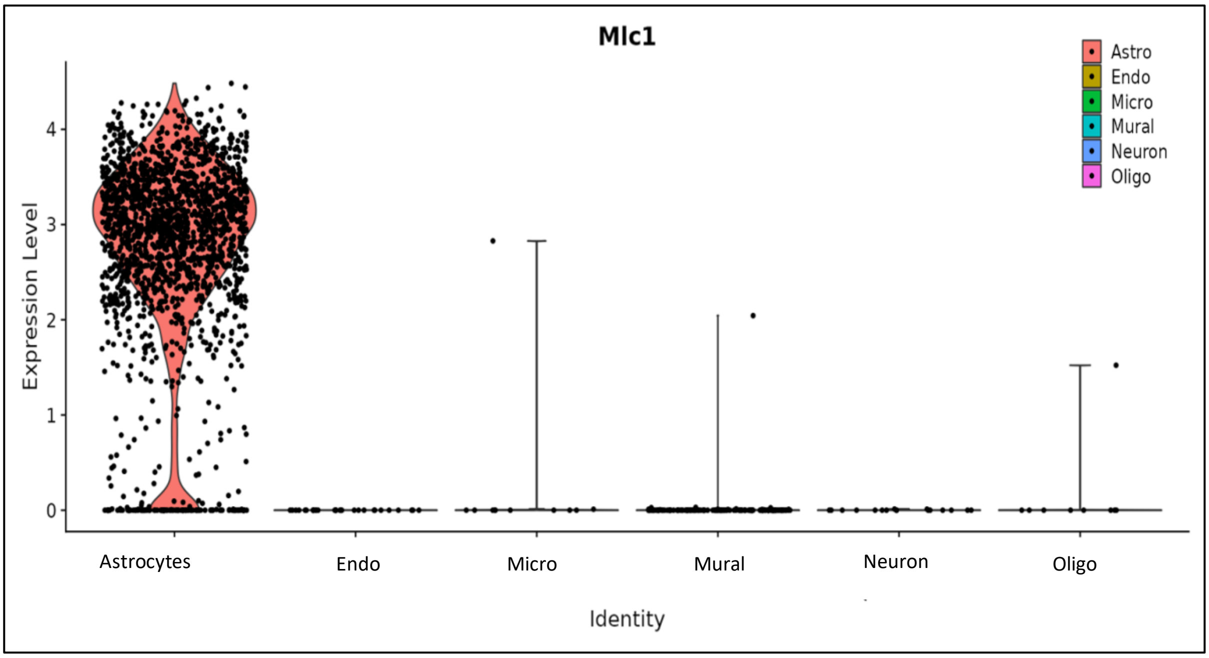

**Figure S1. Astrocytes exhibit nearly exclusive expression of *MLC1* within human brain cells (as curated from: [https://holt-sc.glialab.org/sc/](https://holt-sc.glialab.org/sc/) )**

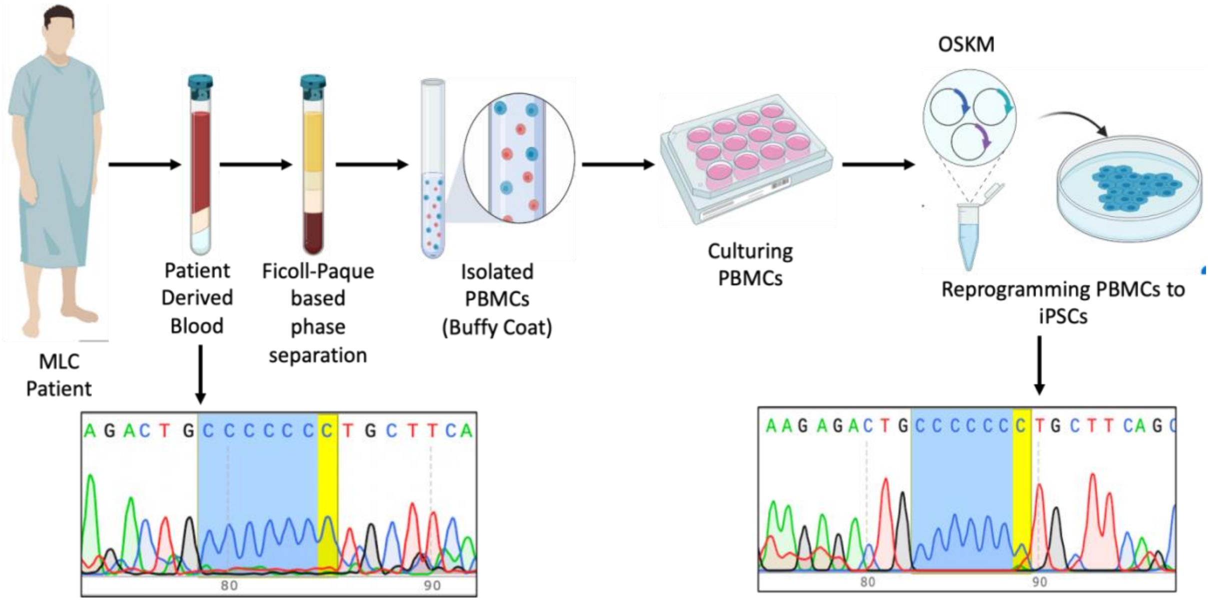

**Figure S2. Sanger sequencing was performed with MLC1 gene amplicon from gDNA isolated from patient-derived blood as well as the cells obtained post-reprogramming.** Sequencing demonstrated that the insertion mutation (136dupC) resulting in MLC is prevalent in the patient-derived blood cells (towards the left side) as well as the reprogrammed cells (towards right side). This estimation helps confirm that the reprogramming procedure didn't affect the integrity of gene locus of our concern, i.e. *MLC1* region which bears the patient mutation.

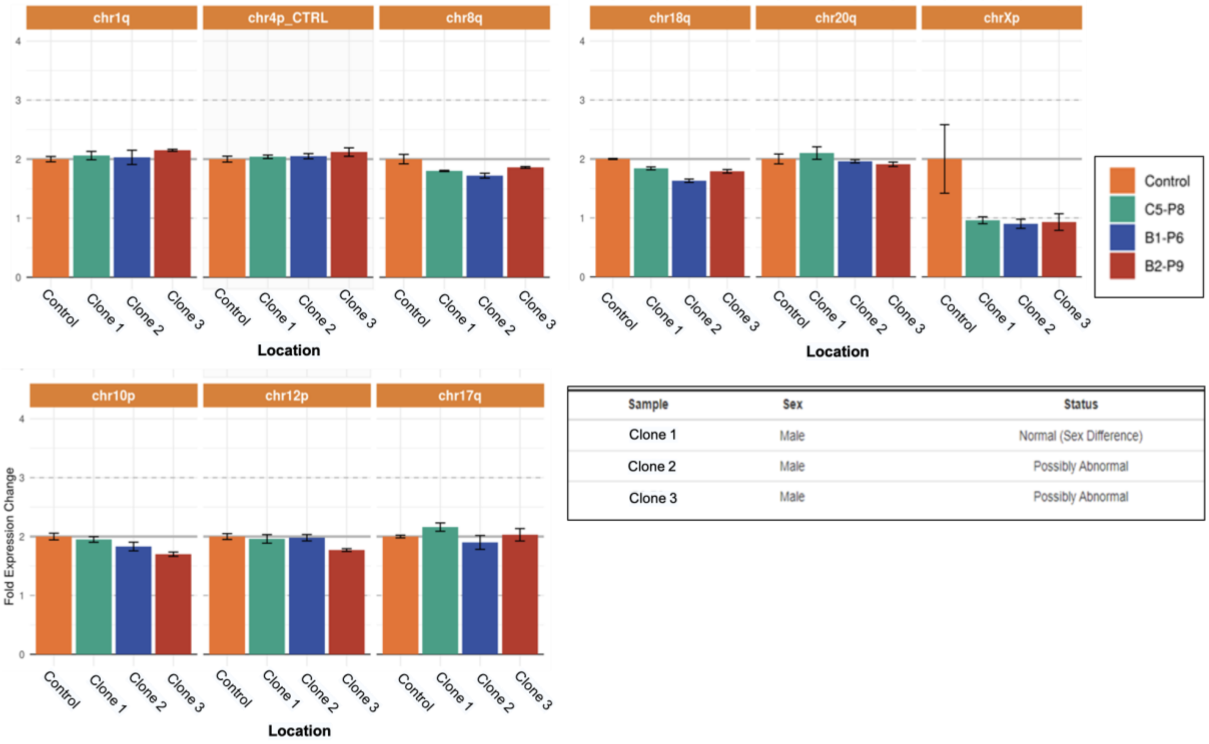

**Figure S3. qPCR based karyotype analysis of genomic DNA extracted from different iPSC clones of MLC patient-derived iPSCs.** Clone 1 bears a normal karyotype with only sex difference as a result of a female genomic DNA control with respect to the male cell line tested.

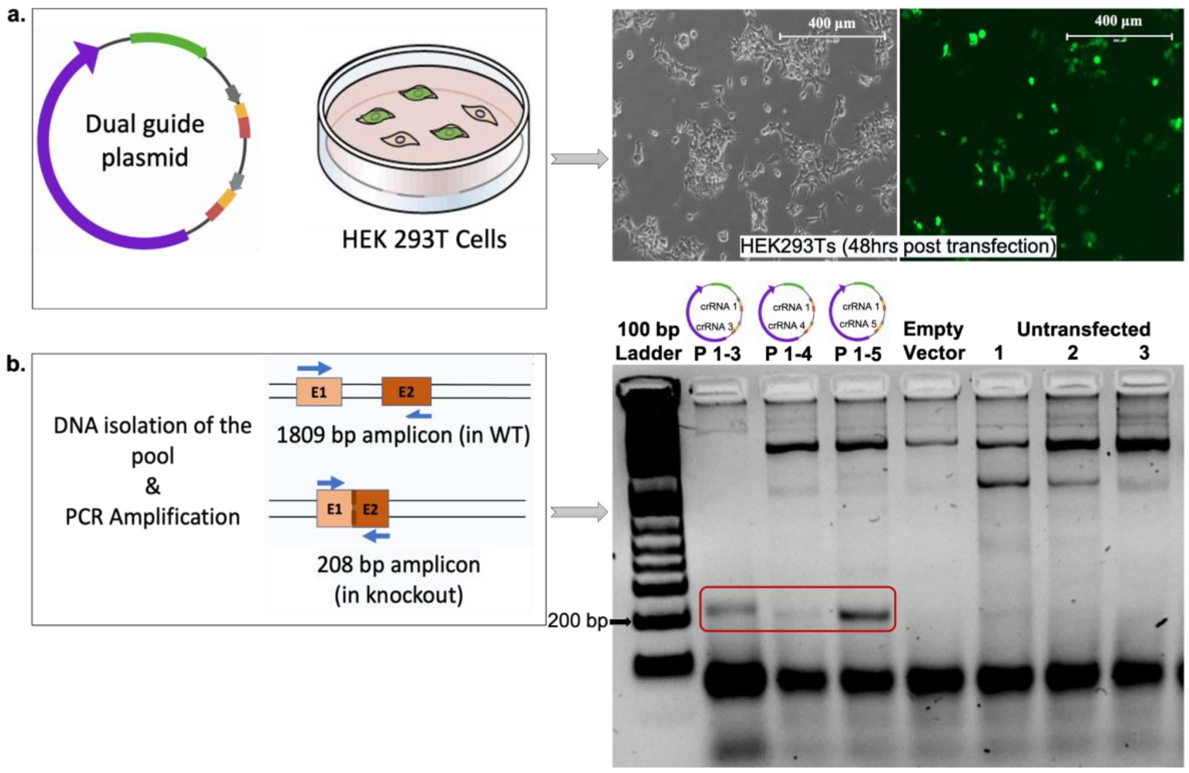

**Figure S4. Transfection efficiency and validation of the dual guide plasmid in HEK293T cells.** a. Fluorescence check of the efficiency of genome editing in the pooled cell population. b. Plasmid combination P1-3 and P1-5 resulted in a knockout. P1-3 was taken forward to perform electroporation in human iPSCs following validation via PCR-based screens in HEK cells' (treated with this dual guide RNA plasmid) genomic DNA.

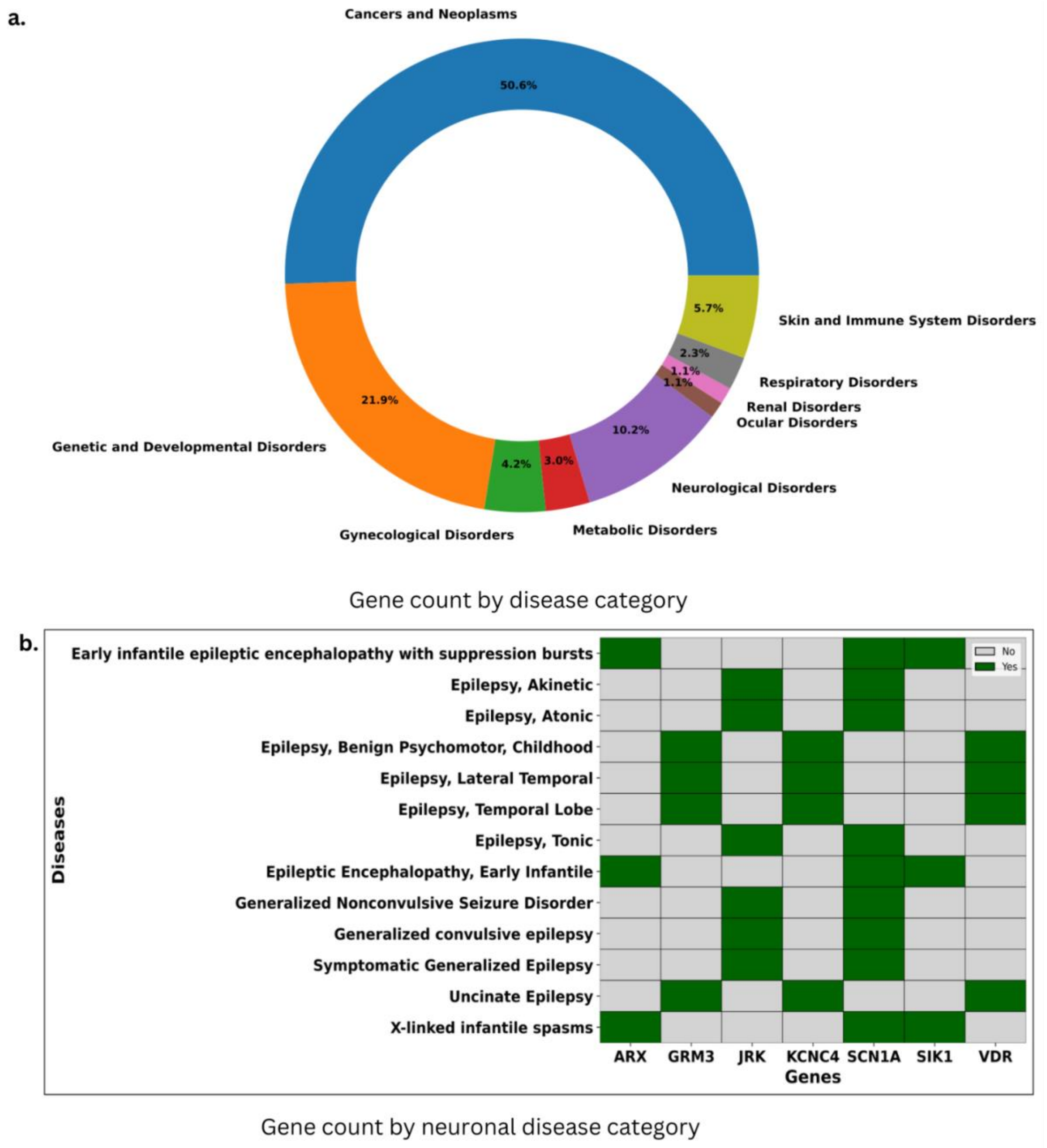

**Figure S5. Association of differentially expressed genes in *MLC1* ablated NSCs with human disorders.** a. System-wide disorders associated with MLC1 NSCs. b. Neurological diseases associated with MLC1 disruption.

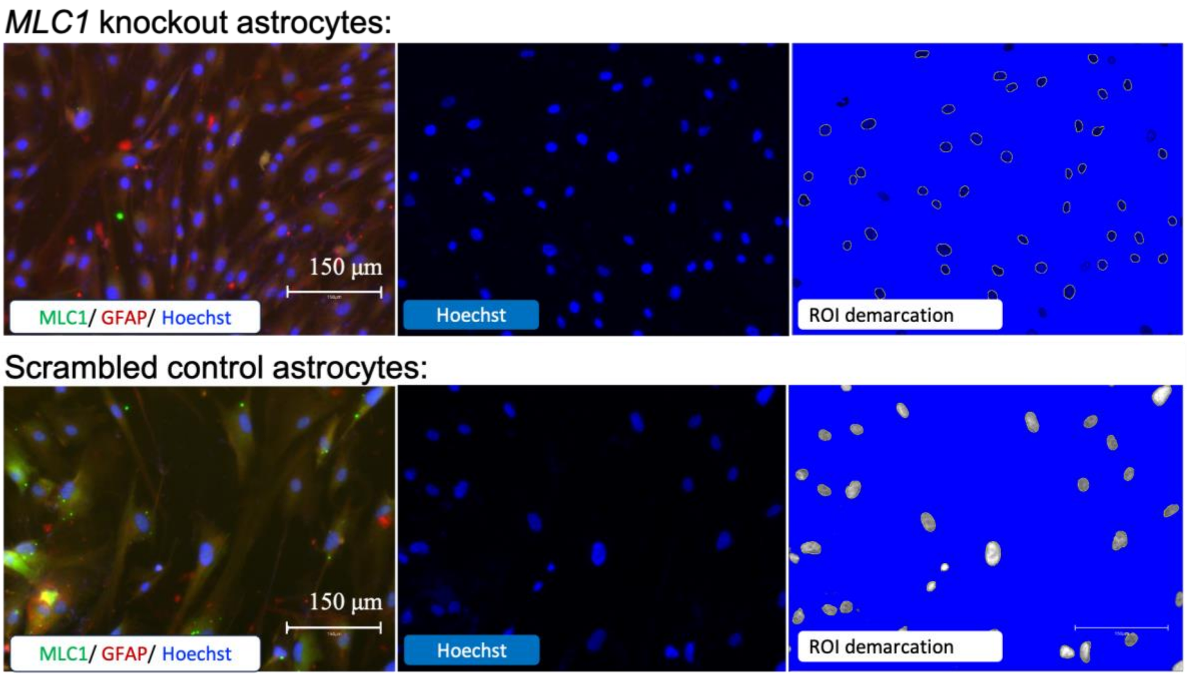

**Figure S6. Estimation of size differences in the size of nuclei of MLC1 knockout astrocytes with respect to Scrambled control astrocytes.** MLC1 knockout astrocytes looked visibly shrunken in size as compared to scrambled control astrocytes.

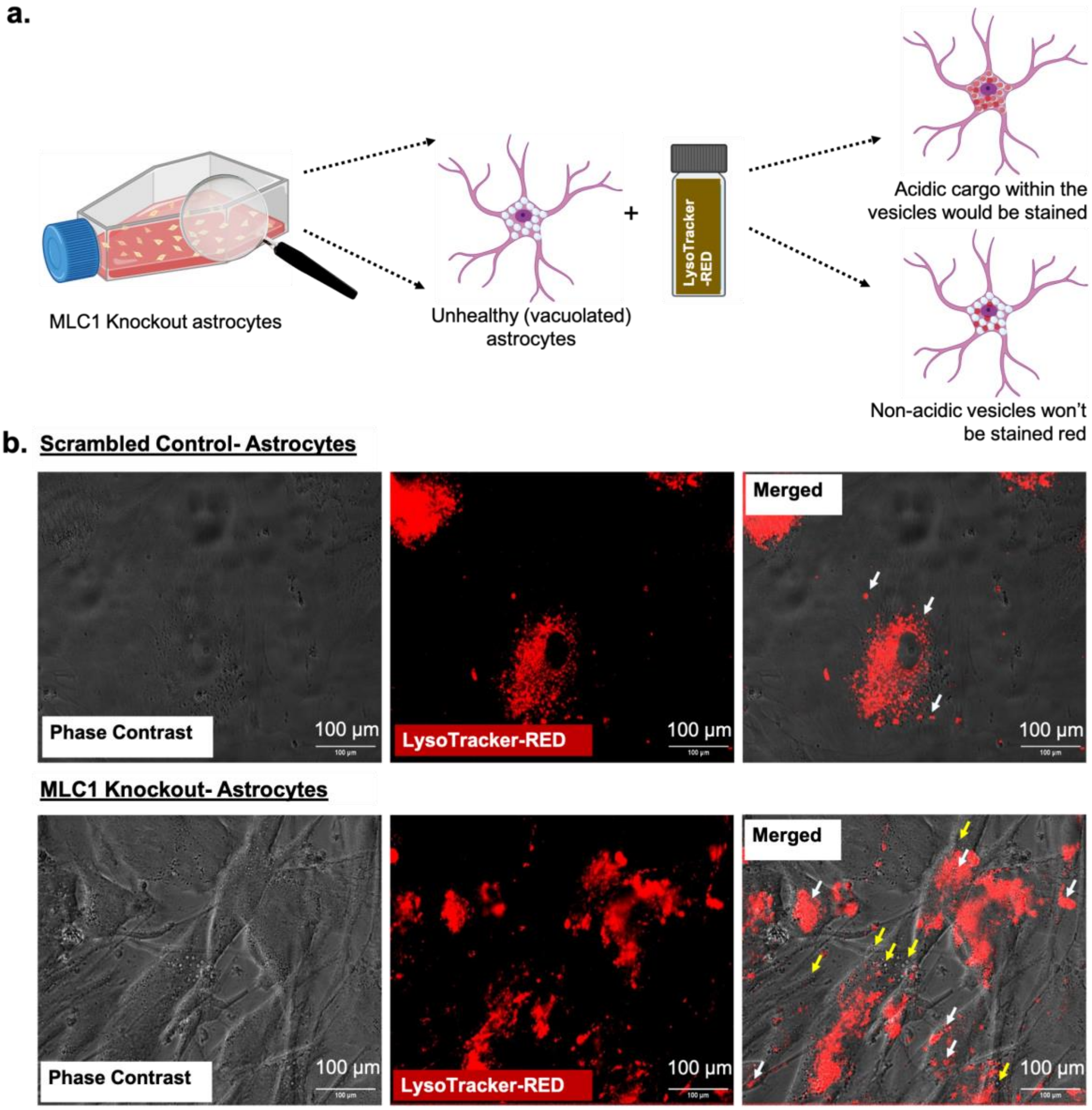

**Figure S7. MLC disease-relevant vacuoles are not associated with lysosomes.** a. Astrocytes derived from MLC1 deleted iPSCs were predicted to develop a vacuolated phenotype representative of MLC disease. b. LysoTracker RED staining was used to observe vacuole acidity. c. LysoTrackerRed did not stain all of the vacuoles observed in the MLC1 gene knockout astrocytes (yellow arrows indicate unstained vacuoles).

**Table S1. List of primers used to develop and validate MLC iPSC model(s).**

| Primer Name | Sequence (5' --> 3') |
|---|---|
| FP-MLC1-271 | GGAGCCATTCAGAGAGGAGC |
| RP-MLC1-271 | GTTGCTCTCTCTGTCCCACC |
| FP-MLC1-196 | AGCCATTCAGAGAGGAGCTG |
| RP-MLC1-196 | CTCCAAGGGCTTAGTGTGGA |
| FP-crRNA1-MLC1 | CACCGCCTGCTTCAGCCACAAGACG |
| RP-crRNA1-MLC1 | AAACCGTCTTGTGGCTGAAGCAGGC |
| FP-crRNA2-MLC1 | CAACCGTCTTGTGGCTGAAGCAGGG |
| RP-crRNA2-MLC1 | AAACCCTGCTTCAGCCACAAGAC |
| FP-crRNA3-MLC1 | CACCGAGGTACAGCGAAAACCCCG |
| RP-crRNA3-MLC1 | AAACCGGGGTTTTCGCTGTACCTC |
| FP-crRNA4-MLC1 | CAC CGA GCC GGG AAC ACG TTC CCC |
| RP-crRNA4-MLC1 | AAACGGGGAACGTGTTCCCGGCTC |
| FP-crRNA5-MLC1 | CACCGTGAAGCTCACAATTGCCGA |
| RP-crRNA5-MLC1 | AAACTCGGCAATTGTGAGCTTCAC |
| FP-MLC1-Genotyping | CGGATGCCCACGCTGGAGCGGGGCCGGCAA |
| RP-MLC1-Genotyping | ACTGGCAGAGGCGTGGAGGAAGCTGCTTAC |
| FP-MLC1donor-RE1 (BamHI) | GCGGCGGGATCC CCCCAGACGCGAAGCC |
| RP-MLC1donor-RE2 (BamHI) | GCGGCGGGATCC CTACTCACCCCCATCA |
| FP-MLC1-196 | AGCCATTCAGAGAGGAGCTG |
| RP-MLC1-196 | CTCCAAGGGCTTAGTGTGGA |
| FP-MLC1-cDNA | ATGACCCAGGAGCCATTCAGAGAG |
| RP-MLC1-cDNA | TCACTGGGCCATTTGCACCACGAC |
| FP-cDNA-MLC-Age1RE | GGCGCCACCGGTATGACCCAGGAGCCATTCAGAGAG |
| RP-cDNA-MLC-Xho1RE | GGCGCCCTCGAGTCACTGGGCCATTTGCACCACGAC |
| Myco2(cb) | CTTCWTCGACTTYCAGACCCAAGGCAT |
| Myco11(cb) | ACACCATGGGAGYTGGTAAT |
| Myco5(cb) | GGTGTGGGTGAGTTATTACAAARTCAATT |
| Myco6(cb) | GGAGTGAGTGGATCCATAAATTGTGA |
| h/m_gDNA_ctrl_FW | TACCAGGTGTCCTCCTCCA |
| h/m_gDNA_ctrl_REV | GATGCCAAAGGCAGCTTTCC |

**Table S2. List of oligos used by qPCR for characterizing the iPSC based MLC model.**

| | |
|---|---|
| TBXT/Brachyury<br>(Primetime std qPCR assay) | 5'-/56-FAM/AGGAGTTCA/ZEN/GCATGATCTGGCCC/3IABKFQ/-3'<br>(For qPCR based Trilineage differentiation assay of MLC1 gene knockout iPSCs) |
| MSI<br>(Primetime std qPCR assay) | 5'-/56-FAM/CGATTGCGC/ZEN/CAGCACTTTATCCAC/3IABKFQ/-3'<br>(For qPCR based Trilineage differentiation assay of MLC1 gene knockout iPSCs) |
| AFP<br>(Primetime std qPCR assay) | 5'-/56-FAM/AATGCTGCA/ZEN/AACTGACCACGCTG/3IABKFQ/-3'<br>(For qPCR based Trilineage differentiation assay of MLC1 gene knockout iPSCs) |
| GAPDH<br>(PrimeTime std qPCR assay) | 5'-/56-FAM/CCGTTGACT/ZEN/CCGACCTTCACCTT/3IABKFQ/-3'<br>(For qPCR based Trilineage differentiation assay of MLC1 gene knockout iPSCs) |

**Table S3. List of reagents used to develop MLC model(s).**

| Assay | Antibody | Dilution | Company, Catalogue No. |
|---|---|---|---|
| Pluripotency Assessment | Anti-Sox2<br>Anti-Oct4<br>Anti-Oct4<br>Anti-Klf4<br>Anti-SSEA4 | 1:100<br>1:100<br>1:500<br>1:100<br>1:400 | Abcam, #ab97959<br>Abcam, #ab19857<br>Cst, #2750<br>Abcam, #ab215036<br>Cst, #MC813 |
| Trilineage Differentiation Efficiency | Anti-Tuj1<br>Anti-SMA<br>Anti-GATA4 | 1:100<br>1:200<br>1:100 | Abcam, #Ab78078<br>Invitrogen, #A25538<br>Abcam, #Ab134057 |
| For Neural Stem Cells | Anti-Sox1<br>Anti-Sox2<br>Anti-Nestin<br>Anti-Pax6 | 1:100<br>1:100<br>1:100<br>1:50 | Abcam, #ab109290<br>Abcam, #ab97959<br>Abcam, #ab6142<br>Abcam, #ab5790 |
| For Astrocytes | Anti-GFAP<br><br>Anti-MLC1 | 1:5000<br>1:1000<br>1:100 | Abcam, #ab7260 (Pd-NSCs)<br>Novus, #NBP1-05198<br>Novus, #NBP1-81555 |
| Secondary Antibodies Used (for Immunofluorescence staining) | Alexa Fluor 488 Goat anti-rabbit<br>Alexa Fluor 488 Goat anti-mouse<br>Alexa Fluor 555 Goat anti-rabbit | 1:500<br>1:500<br>1:500 | Invitrogen, #A11034<br>Invitrogen, #A11001<br>Invitrogen, #A27039 |

| Kits Used | Company, Cat. No. |
|---|---|
| CytoTune-iPS 2.0 Sendai Reprogramming Kit | Invitrogen, #A16517 |
| STEMdiff Trilineage differentiation kit | StemCell Tech, #7550 |
| Wizard® gDNA purification Kit | Promega, #A1125 |
| Trilineage Differentiation media kit | StemCell Tech, #5230 |
| TOPO™ TA Cloning™ Kit | Invitrogen, # K450002 |
| Matrigel (hES Cell Quality) | Corning, #354277 |
| ReLeSR | StemCell Tech, #5872 |
| mTesr1 | StemCell Tech, #85870 |
| mFreSR | StemCell Tech, #05853 |
| StemMACS iPSC brew XF Human | Miltenyi Biotec, #130-104-368 |
| StemMACS cryobrew | Miltenyi Biotec,#130-109-558 |
| Astrocyte Medium | ScienCell, #1801 |
| Trypsin/EDTA Solution, 0.05% | ScienCell, #0183 |
| Luna Universal qPCR MM | NEB, #M30045 |
| StemPRO 34 SFM media & nutrient supplement | Gibco, #10640-019 & #10641-025 |
| Cytokines:<br>Recombinant human IL3<br>Recombinant human IL6<br>Recombinant human SCF<br>Recombinant human FLT3 Ligand | <br>Invitrogen, #PHC0034<br>Gibco, #PHC0065<br>Gibco, #PHC2116<br>Gibco, #PHC9415 |
| DAPI | Sigma, #D9542 |
| Hoechst | Invitrogen, #H21486 |
| L-Glutamine | Gibco, #25030081 |
| LysoTracker RED | Invitrogen, #L7528 |
| Fluorobrite DMEM | Gibco, #A18967-01 |
| Prolong Gold AntiFade with DAPI | Invitrogen, #P36935 |

**Table S4. List of differentially expressed genes found in MLC1 gene ablated NSCs with respect to scrambled control**

| Ensembl gene ID | baseMean | log2FoldChange | lfcSE | stat | p value | p adj | symbol |
|---|---|---|---|---|---|---|---|
| ENSG00000287080 | 1781.982503 | -7.428219595 | 0.8407481201 | -8.835249723 | 9.99E-19 | 3.32E-15 | H3C3 |
| ENSG00000180318 | 44.42711738 | -7.396562147 | 1.951845351 | -3.78952264 | 0.0001509370554 | 0.01229989024 | ALX1 |
| ENSG00000145681 | 415.5791739 | -5.814545843 | 1.098646309 | -5.29246382 | 1.21E-07 | 4.07E-05 | HAPLN1 |
| ENSG00000157554 | 437.7120879 | -5.684924343 | 0.8852190869 | -6.422053509 | 1.34E-10 | 1.31E-07 | ERG |
| ENSG00000135636 | 1237.314936 | -5.320593098 | 1.230648268 | -4.323406805 | 1.54E-05 | 0.002220938294 | DYSF |
| ENSG00000179772 | 293.2215072 | -5.211983785 | 1.348781556 | -3.864216383 | 0.0001114463964 | 0.009982962937 | FOXS1 |
| ENSG00000120149 | 3383.028842 | -5.18810789 | 0.5488416615 | -9.452831762 | 3.30E-21 | 2.74E-17 | MSX2 |
| ENSG00000077274 | 713.6081245 | -5.169353365 | 1.188896744 | -4.348025504 | 1.37E-05 | 0.002038942997 | CAPN6 |
| ENSG00000176971 | 661.6735995 | -4.947184639 | 0.5183233047 | -9.544592331 | 1.37E-21 | 2.27E-17 | FIBIN |
| ENSG00000203805 | 97.7030467 | -4.944365489 | 1.062264592 | -4.654551723 | 3.25E-06 | 0.0006276262208 | PLPP4 |

| | | | | | | | |
|---|---|---|---|---|---|---|---|
| ENSG00000172238 | 335.0436903 | -4.836259979 | 1.304268216 | -3.708025635 | 0.000208881515 | 0.0155715081 | ATOH1 |
| ENSG00000168685 | 511.2481718 | -4.785816879 | 1.354084988 | -3.534354874 | 0.0004087718461 | 0.02471062971 | IL7R |
| ENSG00000077943 | 310.4776653 | -4.774499946 | 0.931389066 | -5.126214297 | 2.96E-07 | 8.47E-05 | ITGA8 |
| ENSG00000019186 | 558.0749085 | -4.750608381 | 1.331024083 | -3.569137811 | 0.0003581579965 | 0.02265412336 | CYP24A1 |
| ENSG00000237515 | 571.2571444 | -4.709177418 | 1.108093376 | -4.249801977 | 2.14E-05 | 0.002845491285 | SHISA9 |
| ENSG00000135333 | 1879.85503 | -4.682571006 | 0.5268842011 | -8.887286801 | 6.26E-19 | 2.60E-15 | EPHA7 |
| ENSG00000078098 | 630.4461844 | -4.657616971 | 0.8324662964 | -5.594961611 | 2.21E-08 | 9.91E-06 | FAP |
| ENSG00000119681 | 10814.47262 | -4.642014289 | 1.202741876 | -3.859526623 | 0.0001136068805 | 0.01004574883 | LTBP2 |
| ENSG00000164188 | 661.4712644 | -4.630576041 | 1.103874505 | -4.194839196 | 2.73E-05 | 0.003491874264 | RANBP3L |
| ENSG00000266524 | 3330.969629 | -4.622209339 | 1.08420425 | -4.263227466 | 2.01E-05 | 0.002745620073 | GDF10 |
| ENSG00000276410 | 55.11335628 | -4.608967979 | 1.068981384 | -4.311551209 | 1.62E-05 | 0.002323250122 | H2BC3 |
| ENSG00000069122 | 200.6916199 | -4.600036571 | 0.8695925524 | -5.289875768 | 1.22E-07 | 4.07E-05 | ADGRF5 |
| ENSG00000164283 | 47.30938724 | -4.576406474 | 1.349268851 | -3.391767674 | 0.0006944329561 | 0.03683309866 | ESM1 |
| ENSG00000125492 | 104.110087 | -4.444746287 | 1.123797341 | -3.955113726 | 7.65E-05 | 0.007578136239 | BARHL1 |
| ENSG00000230838 | 124.5091671 | -4.399639603 | 1.339981277 | -3.283359014 | 0.001025779394 | 0.04725509057 | LINC01614 |
| ENSG00000168329 | 247.4497169 | -4.399612244 | 1.20962877 | -3.637159064 | 0.0002756617101 | 0.01871807289 | CX3CR1 |
| ENSG00000150048 | 188.7569328 | -4.307003796 | 0.7070309108 | -6.0916768 | 1.12E-09 | 6.88E-07 | CLEC1A |
| ENSG00000160801 | 810.6381658 | -4.30596875 | 1.007394623 | -4.274361459 | 1.92E-05 | 0.002700496243 | PTH1R |
| ENSG00000259807 | 127.0295406 | -4.200693384 | 0.985152055 | -4.264005097 | 2.01E-05 | 0.002745620073 | Novel transcript |
| ENSG00000130300 | 86.15382549 | -4.11816671 | 0.9311571773 | -4.422633268 | 9.75E-06 | 0.001558582605 | PLVAP |
| ENSG00000204252 | 59.52940963 | -4.068274982 | 0.8745813815 | -4.651682586 | 3.29E-06 | 0.0006291086847 | HLA-DOA |
| ENSG00000151572 | 1334.279048 | -4.053900059 | 1.104270629 | -3.671111007 | 0.0002414984166 | 0.01693953451 | ANO4 |
| ENSG00000147889 | 354.5555545 | -4.048593815 | 1.221374118 | -3.314785989 | 0.0009171331049 | 0.04429362938 | CDKN2A |
| ENSG00000104415 | 1729.479005 | -4.039071416 | 1.095627109 | -3.686538407 | 0.0002273251472 | 0.01635953787 | CCN4 |
| ENSG00000274618 | 1416.736091 | -4.036112728 | 0.7233158964 | -5.580013861 | 2.40E-08 | 1.05E-05 | H4C6 |
| ENSG00000145824 | 1025.365808 | -3.989381062 | 0.9894552708 | -4.031896317 | 5.53E-05 | 0.005784796235 | CXCL14 |
| ENSG00000171992 | 4912.0877 | -3.981679448 | 0.5951282167 | -6.690456505 | 2.22E-11 | 2.84E-08 | SYNPO |
| ENSG00000181541 | 1392.368155 | -3.972780315 | 0.8993566205 | -4.41735817 | 9.99E-06 | 0.001581885843 | MAB21L2 |
| ENSG00000115232 | 2120.650458 | -3.964413833 | 0.5891392586 | -6.729162545 | 1.71E-11 | 2.36E-08 | ITGA4 |
| ENSG00000198848 | 104.6812476 | -3.942352761 | 1.066442217 | -3.696733585 | 0.0002183913165 | 0.0160643241 | CES1 |
| ENSG00000123572 | 459.2765469 | -3.934445994 | 0.8805754792 | -4.468039466 | 7.89E-06 | 0.001299301295 | NRK |
| ENSG00000183542 | 35.06657195 | -3.914717009 | 1.126311301 | -3.475697177 | 0.0005095272757 | 0.02930927831 | KLRC4 |
| ENSG00000107562 | 2533.775534 | -3.886272153 | 0.798253608 | -4.868468008 | 1.12E-06 | 0.0002709633103 | CXCL12 |
| ENSG00000234779 | 69.63657483 | -3.868108185 | 0.9133538797 | -4.235059675 | 2.28E-05 | 0.002967527114 | BNC2-AS1 |

| | | | | | | | |
|---|---|---|---|---|---|---|---|
| ENSG00000113361 | 16033.82928 | -3.867969839 | 0.8082676392 | -4.785506251 | 1.71E-06 | 0.0003682256543 | CDH6 |
| ENSG00000170381 | 342.9697438 | -3.864134356 | 0.827077647 | -4.672033357 | 2.98E-06 | 0.0005832726157 | SEMA3E |
| ENSG00000116690 | 99.67845277 | -3.85949949 | 1.095249417 | -3.523854413 | 0.0004253178717 | 0.02516186583 | PRG4 |
| ENSG00000115461 | 77859.26093 | -3.855980692 | 1.045582627 | -3.687877546 | 0.0002261324227 | 0.01634445824 | IGFBP5 |
| ENSG00000135406 | 93.85721217 | -3.850827845 | 1.071350616 | -3.594367508 | 0.0003251805682 | 0.02128268412 | PRPH |
| ENSG00000174130 | 304.8962093 | -3.836162752 | 0.5760744956 | -6.659143533 | 2.75E-11 | 3.27E-08 | TLR6 |
| ENSG00000122641 | 3738.807781 | -3.810981221 | 0.9873884948 | -3.859657309 | 0.0001135461444 | 0.01004574883 | INHBA |
| ENSG00000133110 | 3975.676462 | -3.80085238 | 0.6495940656 | -5.851119309 | 4.88E-09 | 2.62E-06 | POSTN |
| ENSG00000102924 | 161.8771382 | -3.767545755 | 0.9286979187 | -4.056804349 | 4.97E-05 | 0.005335630718 | CBLN1 |
| ENSG00000052850 | 575.154196 | -3.76387697 | 0.9812636442 | -3.835744851 | 0.0001251843187 | 0.0108388756 | ALX4 |
| ENSG00000069535 | 33.97744577 | -3.761678146 | 1.132002904 | -3.323028706 | 0.0008904575058 | 0.04341045624 | MAOB |
| ENSG00000137203 | 1353.411109 | -3.661739277 | 0.9705158102 | -3.772982612 | 0.0001613075337 | 0.01295447556 | TFAP2A |
| ENSG00000144057 | 4017.914849 | -3.600692551 | 0.590529857 | -6.09739289 | 1.08E-09 | 6.88E-07 | ST6GAL2 |
| ENSG00000106034 | 2202.285612 | -3.600544757 | 0.9416392295 | -3.823698763 | 0.0001314645193 | 0.0113236589 | CPED1 |
| ENSG00000164161 | 1110.94746 | -3.59429979 | 1.034754052 | -3.473578848 | 0.0005135664432 | 0.02939542644 | HHIP |
| ENSG00000086570 | 244.6581063 | -3.594277671 | 0.8247881245 | -4.357819376 | 1.31E-05 | 0.00196739385 | FAT2 |
| ENSG00000170577 | 103.1584185 | -3.58926992 | 0.8608145001 | -4.169620656 | 3.05E-05 | 0.00375711089 | SIX2 |
| ENSG00000178031 | 2332.371754 | -3.513688458 | 0.893780541 | -3.931265335 | 8.45E-05 | 0.008027012413 | ADAMTSL1 |
| ENSG00000234362 | 52.23682399 | -3.460963474 | 1.037370344 | -3.336285345 | 0.0008490596824 | 0.04188358504 | LINC01914 |
| ENSG00000274441 | 41.89915292 | -3.451579024 | 0.9721582377 | -3.550429231 | 0.0003846035237 | 0.02383490832 | Novel transcript |
| ENSG00000166923 | 496.4144239 | -3.413383155 | 1.01020761 | -3.378892736 | 0.0007277839802 | 0.03754454534 | GREM1 |
| ENSG00000085276 | 259.7659379 | -3.392397334 | 0.9529773597 | -3.559787963 | 0.0003711543503 | 0.02328328271 | MECOM |
| ENSG00000164616 | 71.33939242 | -3.3778624 | 0.843903793 | -4.002662896 | 6.26E-05 | 0.006467202208 | FBXL21P |
| ENSG00000140937 | 7724.272033 | -3.371720746 | 0.5972338202 | -5.645562311 | 1.65E-08 | 7.82E-06 | CDH11 |
| ENSG00000103196 | 528.6299155 | -3.368473905 | 0.7423744536 | -4.537432409 | 5.69E-06 | 0.001021942769 | CRISPLD2 |
| ENSG00000171812 | 496.8013751 | -3.365907815 | 0.6559453311 | -5.131384669 | 2.88E-07 | 8.47E-05 | COL8A2 |
| ENSG00000278709 | 125.6417432 | -3.347569579 | 0.9324824617 | -3.589954467 | 0.000330735729 | 0.0214204041 | NKILA |
| ENSG00000101825 | 10992.53846 | -3.346396621 | 0.941096043 | -3.555850272 | 0.0003767586127 | 0.02345780965 | MXRA5 |
| ENSG00000129009 | 4771.41637 | -3.340332041 | 0.6715876065 | -4.973784519 | 6.57E-07 | 0.0001705474079 | ISLR |
| ENSG00000137809 | 8693.748955 | -3.330901443 | 1.005055403 | -3.314147093 | 0.0009192313604 | 0.04429362938 | ITGA11 |
| ENSG00000112964 | 437.4092015 | -3.317802588 | 0.5543912193 | -5.984587188 | 2.17E-09 | 1.29E-06 | GHR |
| ENSG00000155011 | 2282.311974 | -3.308675217 | 0.7775356552 | -4.255335681 | 2.09E-05 | 0.002814660922 | DKK2 |
| ENSG00000234840 | 86.64630293 | -3.307427154 | 1.003113306 | -3.29716208 | 0.0009766712238 | 0.0458649221 | LINC01239 |
| ENSG00000104313 | 1276.805281 | -3.298023159 | 0.6707418007 | -4.916978717 | 8.79E-07 | 0.000221376525 | EYA1 |

| | | | | | | | |
|---|---|---|---|---|---|---|---|
| ENSG00000134853 | 2525.357911 | -3.260241622 | 0.9885144527 | -3.298122362 | 0.0009733370836 | 0.04583783478 | PDGFRA |
| ENSG00000106991 | 3473.853478 | -3.256010921 | 0.5327735161 | -6.11143539 | 9.87E-10 | 6.57E-07 | ENG |
| ENSG00000119699 | 1422.632317 | -3.232147781 | 0.6400149708 | -5.050112777 | 4.42E-07 | 0.0001183921881 | TGFB3 |
| ENSG00000137959 | 1618.072342 | -3.221444053 | 0.5191660625 | -6.205035895 | 5.47E-10 | 3.95E-07 | IFI44L |
| ENSG00000170340 | 1740.848137 | -3.18473079 | 0.9648123059 | -3.300881188 | 0.0009638168822 | 0.04564812493 | B3GNT2 |
| ENSG00000173068 | 3824.718431 | -3.174094898 | 0.6725976097 | -4.719158754 | 2.37E-06 | 0.00048011319 | BNC2 |
| ENSG00000137507 | 172.3686344 | -3.155971005 | 0.8899803611 | -3.546113086 | 0.0003909583932 | 0.02389445709 | LRRC32 |
| ENSG00000137965 | 206.9764611 | -3.145224144 | 0.5701504636 | -5.516480903 | 3.46E-08 | 1.44E-05 | IFI44 |
| ENSG00000115594 | 885.0157989 | -3.11694622 | 0.4938526971 | -6.311489719 | 2.76E-10 | 2.42E-07 | IL1R1 |
| ENSG00000169174 | 448.6055699 | -3.11190457 | 0.9169677525 | -3.393690304 | 0.0006895762784 | 0.03683309866 | PCSK9 |
| ENSG00000172986 | 1798.872451 | -3.091236414 | 0.6328606778 | -4.884544927 | 1.04E-06 | 0.0002572204317 | GXYLT2 |
| ENSG00000149380 | 93.62257829 | -3.081857717 | 0.875084448 | -3.521783211 | 0.000428654527 | 0.02526933637 | P4HA3 |
| ENSG00000116774 | 1422.103926 | -3.079903692 | 0.5158345858 | -5.970719639 | 2.36E-09 | 1.35E-06 | OLFML3 |
| ENSG00000089327 | 403.4000926 | -3.05358044 | 0.5718254425 | -5.34005697 | 9.29E-08 | 3.43E-05 | FXYD5 |
| ENSG00000277639 | 413.757847 | -3.022806301 | 0.5908853162 | -5.115724182 | 3.13E-07 | 8.81E-05 | CHD9NB |
| ENSG00000181234 | 1103.562168 | -3.011074434 | 0.4769967033 | -6.312568648 | 2.74E-10 | 2.42E-07 | TMEM132C |
| ENSG00000124225 | 3165.721404 | -2.994369844 | 0.8842392041 | -3.386379873 | 0.0007082127506 | 0.03725736951 | PMEPA1 |
| ENSG00000177409 | 234.1729826 | -2.965045212 | 0.6464988035 | -4.586311987 | 4.51E-06 | 0.0008241563464 | SAMD9L |
| ENSG00000183160 | 370.0144369 | -2.955677496 | 0.6539304597 | -4.519865151 | 6.19E-06 | 0.001082817945 | TMEM119 |
| ENSG00000082497 | 3829.794182 | -2.920490257 | 0.5341622012 | -5.467422162 | 4.57E-08 | 1.85E-05 | SERTAD4 |
| ENSG00000156427 | 136.1811775 | -2.914718568 | 0.7566812158 | -3.851976906 | 0.0001171680754 | 0.01030583114 | FGF18 |
| ENSG00000084636 | 1268.724348 | -2.906820327 | 0.4631632246 | -6.27601712 | 3.47E-10 | 2.89E-07 | COL16A1 |
| ENSG00000250722 | 355.5151257 | -2.901328141 | 0.6080263212 | -4.77171471 | 1.83E-06 | 0.000389308753 | SELENOP |
| ENSG00000147883 | 1691.755844 | -2.894158997 | 0.7933073248 | -3.648219179 | 0.0002640643228 | 0.01814586378 | CDKN2B |
| ENSG00000106714 | 1377.014434 | -2.887736256 | 0.5812752772 | -4.967932355 | 6.77E-07 | 0.0001730700563 | CNTNAP3 |
| ENSG00000166250 | 3499.559762 | -2.882163066 | 0.5607579499 | -5.139763185 | 2.75E-07 | 8.31E-05 | CLMP |
| ENSG00000162878 | 5864.310691 | -2.874912704 | 0.5933852648 | -4.844934438 | 1.27E-06 | 0.0002884229722 | PKDCC |
| ENSG00000049089 | 554.5137572 | -2.872916359 | 0.6079865125 | -4.725296204 | 2.30E-06 | 0.0004715883506 | COL9A2 |
| ENSG00000105976 | 150.3832191 | -2.870888181 | 0.6517258025 | -4.40505527 | 1.06E-05 | 0.001658587614 | MET |
| ENSG00000108691 | 1528.438481 | -2.868564592 | 0.8385589176 | -3.42082653 | 0.0006243114001 | 0.03402804169 | CCL2 |
| ENSG00000120708 | 25190.7659 | -2.867353959 | 0.6874953353 | -4.170724966 | 3.04E-05 | 0.00375711089 | TGFBI |
| ENSG00000170891 | 339.2936459 | -2.828759948 | 0.6006954975 | -4.709141254 | 2.49E-06 | 0.0004982445866 | CYTL1 |
| ENSG00000171621 | 697.5513935 | -2.826600009 | 0.8557188916 | -3.303187573 | 0.0009559242312 | 0.04553376624 | SPSB1 |
| ENSG00000164318 | 1452.41382 | -2.818381407 | 0.646302227 | -4.360779353 | 1.30E-05 | 0.00196739385 | EGFLAM |

| | | | | | | | |
|---|---|---|---|---|---|---|---|
| ENSG00000177468 | 122.770758 | -2.80999455 | 0.8491575318 | -3.309155775 | 0.0009357775656 | 0.04483102666 | OLIG3 |
| ENSG00000116194 | 419.4489582 | -2.803008893 | 0.5340488865 | -5.248599826 | 1.53E-07 | 4.81E-05 | ANGPTL1 |
| ENSG00000153246 | 484.3186754 | -2.775400611 | 0.7569542869 | -3.666536618 | 0.0002458577267 | 0.01710099937 | PLA2R1 |
| ENSG00000162882 | 124.6375512 | -2.757417172 | 0.7136781153 | -3.863670628 | 0.0001116958076 | 0.009982962937 | HAAO |
| ENSG00000205562 | 175.8324398 | -2.757074642 | 0.6757472146 | -4.080038485 | 4.50E-05 | 0.004974338738 | FLRT2-AS1 |
| ENSG00000174125 | 51.6928918 | -2.742728832 | 0.8345314868 | -3.28654925 | 0.001014230368 | 0.04709678546 | TLR1 |
| ENSG00000126785 | 3147.358239 | -2.732674135 | 0.5637995032 | -4.84688993 | 1.25E-06 | 0.0002884229722 | RHOJ |
| ENSG00000279118 | 238.5205292 | -2.716528105 | 0.5617173698 | -4.836111986 | 1.32E-06 | 0.0002974423309 | |
| ENSG00000173157 | 142.9748146 | -2.679177458 | 0.7243414495 | -3.698776951 | 0.0002166408753 | 0.01600639071 | ADAMTS20 |
| ENSG00000260428 | 215.6187726 | -2.638484609 | 0.7917777699 | -3.332354998 | 0.0008611433247 | 0.04230728337 | SCX |
| ENSG00000159167 | 1004.752104 | -2.630537142 | 0.665816953 | -3.950841338 | 7.79E-05 | 0.007594696 | STC1 |
| ENSG00000225614 | 1125.237777 | -2.619790257 | 0.6439738672 | -4.068162375 | 4.74E-05 | 0.005148589425 | ZNF469 |
| ENSG00000103489 | 4212.798013 | -2.614005821 | 0.7003454065 | -3.732452297 | 0.0001896246162 | 0.01446018174 | XYLT1 |
| ENSG00000111424 | 189.5077894 | -2.612727896 | 0.6316304192 | -4.136482058 | 3.53E-05 | 0.004267932261 | VDR |
| ENSG00000113209 | 307.3487762 | -2.595898145 | 0.7690012859 | -3.375674648 | 0.0007363494022 | 0.03778108785 | PCDHB5 |
| ENSG00000230630 | 1160.298331 | -2.580944715 | 0.7156750767 | -3.606307945 | 0.0003105845353 | 0.02048871951 | DNM3OS |
| ENSG00000119630 | 347.6659411 | -2.553725679 | 0.6591490563 | -3.87427647 | 0.000106941951 | 0.009822115989 | PGF |
| ENSG00000143341 | 8919.175655 | -2.539575523 | 0.5987933862 | -4.241154931 | 2.22E-05 | 0.00291080436 | HMCN1 |
| ENSG00000165424 | 4520.515872 | -2.513484263 | 0.621314665 | -4.045428838 | 5.22E-05 | 0.00553011966 | ZCCHC24 |
| ENSG00000145569 | 118.0005706 | -2.501490396 | 0.6775837211 | -3.69178054 | 0.0002226895822 | 0.01623680533 | OTULINL |
| ENSG00000157227 | 11406.74097 | -2.492987371 | 0.7636481506 | -3.264575929 | 0.001096280754 | 0.04979391053 | MMP14 |
| ENSG00000276386 | 270.5800532 | -2.470975264 | 0.625341156 | -3.951403551 | 7.77E-05 | 0.007594696 | CNTNAP3P2 |
| ENSG00000173705 | 210.8592803 | -2.448713172 | 0.6939144412 | -3.528840195 | 0.0004173850933 | 0.02504913282 | SUSD5 |
| ENSG00000286379 | 197.9501891 | -2.419292961 | 0.5898134124 | -4.101793738 | 4.10E-05 | 0.004700116665 | |
| ENSG00000142178 | 380.4409727 | -2.413286669 | 0.7243105151 | -3.331839892 | 0.0008627387549 | 0.04230728337 | SIK1 |
| ENSG00000130052 | 449.4359682 | -2.398574721 | 0.628581116 | -3.815855521 | 0.0001357118997 | 0.01134605267 | STARD8 |
| ENSG00000254535 | 261.2621458 | -2.386479225 | 0.5792894751 | -4.119666121 | 3.79E-05 | 0.004473409838 | PABPC4L |
| ENSG00000099139 | 580.0942022 | -2.383866502 | 0.5254752697 | -4.536591233 | 5.72E-06 | 0.001021942769 | PCSK5 |
| ENSG00000028116 | 123.3362985 | -2.373771808 | 0.6732436327 | -3.525873387 | 0.0004220887094 | 0.02511986917 | VRK2 |
| ENSG00000143127 | 466.6424062 | -2.370920372 | 0.7047322222 | -3.364285465 | 0.0007674212169 | 0.03899355786 | ITGA10 |
| ENSG00000077238 | 1047.086031 | -2.339114794 | 0.713334189 | -3.279128955 | 0.001041280416 | 0.04781835809 | IL4R |
| ENSG00000102287 | 276.9418647 | -2.339002486 | 0.5658804103 | -4.133386566 | 3.57E-05 | 0.004267932261 | GABRE |
| ENSG00000146122 | 1059.902905 | -2.326246003 | 0.4636421095 | -5.017331161 | 5.24E-07 | 0.0001382541855 | DAAM2 |
| ENSG00000178695 | 4825.710055 | -2.318002489 | 0.5119310449 | -4.527958428 | 5.96E-06 | 0.001053259406 | KCTD12 |

| | | | | | | | |
|---|---|---|---|---|---|---|---|
| ENSG00000205413 | 243.7751217 | -2.266916688 | 0.6568750396 | -3.451062305 | 0.0005583846494 | 0.03083915752 | SAMD9 |
| ENSG00000179403 | 931.3157817 | -2.258290877 | 0.6671203741 | -3.385132526 | 0.000711438969 | 0.03730902657 | VWA1 |
| ENSG00000169851 | 3951.573771 | -2.257758787 | 0.5019601605 | -4.497884423 | 6.86E-06 | 0.001188494579 | PCDH7 |
| ENSG00000128606 | 1591.377709 | -2.254983481 | 0.553847428 | -4.071488586 | 4.67E-05 | 0.005108998335 | LRRC17 |
| ENSG00000121297 | 1182.406025 | -2.250255265 | 0.5244494716 | -4.290699841 | 1.78E-05 | 0.002530697404 | TSHZ3 |
| ENSG00000139263 | 2113.845897 | -2.247913162 | 0.6718590138 | -3.345810826 | 0.000820423483 | 0.04108048187 | LRIG3 |
| ENSG00000185070 | 7701.239086 | -2.233514598 | 0.5031411063 | -4.439141565 | 9.03E-06 | 0.001457721384 | FLRT2 |
| ENSG00000050555 | 2107.170962 | -2.214950089 | 0.6091114983 | -3.636362301 | 0.0002765153488 | 0.01871807289 | LAMC3 |
| ENSG00000165617 | 1072.153133 | -2.157528514 | 0.6350851445 | -3.397227179 | 0.0006807242822 | 0.03650438861 | DACT1 |
| ENSG00000158270 | 2266.074773 | -2.155864884 | 0.6243396902 | -3.453031928 | 0.0005543233889 | 0.03083915752 | COLEC12 |
| ENSG00000008516 | 927.4310831 | -2.153025128 | 0.5928030747 | -3.63193988 | 0.0002812986647 | 0.01893242511 | MMP25 |
| ENSG00000176692 | 389.791061 | -2.149120555 | 0.526656486 | -4.080687531 | 4.49E-05 | 0.004974338738 | FOXC2 |
| ENSG00000123104 | 940.5246483 | -2.121236178 | 0.5658888298 | -3.748503357 | 0.0001778929569 | 0.0138407525 | ITPR2 |
| ENSG00000101665 | 963.0535277 | -2.117926862 | 0.5363612314 | -3.948694905 | 7.86E-05 | 0.007594696 | SMAD7 |
| ENSG00000007944 | 1768.864088 | -2.095877448 | 0.4940336903 | -4.242377573 | 2.21E-05 | 0.00291080436 | MYLIP |
| ENSG00000144285 | 482.3034957 | -2.085560031 | 0.6049139482 | -3.447697044 | 0.0005653878306 | 0.03101982606 | SCN1A |
| ENSG00000204217 | 8302.263458 | -2.081198318 | 0.6372793584 | -3.265755105 | 0.001091726201 | 0.04972289417 | BMPR2 |
| ENSG00000275126 | 288.3421225 | -2.063436962 | 0.6297979378 | -3.276347599 | 0.001051590491 | 0.04815878877 | H4C13 |
| ENSG00000120820 | 730.1649161 | -2.058434104 | 0.5560130956 | -3.702132415 | 0.000213794981 | 0.0158666418 | GLT8D2 |
| ENSG00000111799 | 42581.90988 | -2.04348118 | 0.6195808711 | -3.298166995 | 0.0009731823744 | 0.04583783478 | COL12A1 |
| ENSG00000127863 | 3153.347758 | -2.040601691 | 0.5389048141 | -3.78657165 | 0.0001527400678 | 0.01238610189 | TNFRSF19 |
| ENSG00000126778 | 855.8168355 | -2.036760252 | 0.4991427045 | -4.080516922 | 4.49E-05 | 0.004974338738 | SIX1 |
| ENSG00000187068 | 1017.006173 | -2.023662207 | 0.5218126096 | -3.878139719 | 0.0001052582631 | 0.009744321367 | C3orf70 |
| ENSG00000174749 | 205.6353802 | -2.005081776 | 0.6046006548 | -3.316373808 | 0.0009119376157 | 0.04419839919 | FAM241A |
| ENSG00000088756 | 2807.715325 | -1.994933229 | 0.5744022744 | -3.473059419 | 0.0005145614229 | 0.02939542644 | ARHGAP28 |
| ENSG00000122870 | 2708.968355 | -1.979034397 | 0.4984226107 | -3.970595142 | 7.17E-05 | 0.007311838844 | BICC1 |
| ENSG00000182022 | 2091.057638 | -1.957120509 | 0.4721260615 | -4.145334622 | 3.39E-05 | 0.004147661405 | CHST15 |
| ENSG00000124813 | 2810.702286 | -1.955193021 | 0.5578218929 | -3.505048916 | 0.000456523397 | 0.02662892966 | RUNX2 |
| ENSG00000182326 | 331.9165362 | -1.939252963 | 0.560060031 | -3.462580538 | 0.0005350217665 | 0.03025238723 | C1S |
| ENSG00000160963 | 4844.890738 | -1.9333276 | 0.5687698895 | -3.399138449 | 0.0006759848839 | 0.03636754923 | COL26A1 |
| ENSG00000100599 | 1354.890493 | -1.862861563 | 0.5131855822 | -3.629995906 | 0.0002834257151 | 0.01899866568 | RIN3 |
| ENSG00000263528 | 261.804931 | -1.859676095 | 0.5561421497 | -3.343886264 | 0.0008261359901 | 0.0412422964 | IKBKE |
| ENSG00000131398 | 996.2318239 | -1.849670808 | 0.5629350856 | -3.285762169 | 0.001017068454 | 0.04709678546 | KCNC3 |
| ENSG00000200488 | 1485.568635 | -1.815932361 | 0.4977704987 | -3.648131751 | 0.0002641541768 | 0.01814586378 | RN7SKP203 |

| | | | | | | | |
|---|---|---|---|---|---|---|---|
| ENSG00000154654 | 454.2235419 | -1.812150348 | 0.5306633895 | -3.414877272 | 0.0006381075599 | 0.03466634012 | NCAM2 |
| ENSG00000198715 | 423.4747336 | -1.783828566 | 0.5162180337 | -3.455571967 | 0.0005491265769 | 0.03073629702 | GLMP |
| ENSG00000111145 | 2491.499103 | -1.758069689 | 0.5093939659 | -3.451296652 | 0.0005578999893 | 0.03083915752 | ELK3 |
| ENSG00000112559 | 3708.434227 | -1.738908505 | 0.5140624932 | -3.382679204 | 0.0007178242817 | 0.03740787103 | MDFI |
| ENSG00000162733 | 4915.740486 | -1.738227646 | 0.5248674467 | -3.31174596 | 0.0009271569758 | 0.04454640915 | DDR2 |
| ENSG00000125354 | 3323.409304 | -1.737666874 | 0.5025575731 | -3.457647376 | 0.0005449141075 | 0.03070729533 | SEPTIN6 |
| ENSG00000197380 | 1814.314898 | -1.719356729 | 0.4700740406 | -3.657629608 | 0.0002545585458 | 0.01763242194 | DACT3 |
| ENSG00000198121 | 3280.993554 | -1.716560827 | 0.5191152901 | -3.306704426 | 0.0009440044237 | 0.04509519982 | LPAR1 |
| ENSG00000124212 | 3008.407394 | -1.714282219 | 0.5212278215 | -3.288930767 | 0.001005687583 | 0.04696222015 | PTGIS |
| ENSG00000082397 | 1231.751952 | -1.694044567 | 0.4707698559 | -3.598455904 | 0.0003201121126 | 0.0210337698 | EPB41L3 |
| ENSG00000153071 | 5565.023025 | -1.692101208 | 0.4710816448 | -3.591948927 | 0.0003282141839 | 0.02139699056 | DAB2 |
| ENSG00000104447 | 3339.509209 | -1.689608887 | 0.4751581322 | -3.555887551 | 0.000376705187 | 0.02345780965 | TRPS1 |
| ENSG00000185483 | 1405.634147 | -1.670216114 | 0.496557863 | -3.36358809 | 0.0007693627873 | 0.03899355786 | ROR1 |
| ENSG00000144724 | 5316.220856 | -1.66415551 | 0.4511744336 | -3.688496923 | 0.0002255827541 | 0.01634445824 | PTPRG |
| ENSG00000184232 | 3141.385117 | -1.659937557 | 0.4521596233 | -3.671131767 | 0.000241478799 | 0.01693953451 | OAF |
| ENSG00000155090 | 1747.898668 | -1.531022336 | 0.460434846 | -3.325166088 | 0.0008836587766 | 0.04320571618 | KLF10 |
| ENSG00000139926 | 4142.420117 | -1.521754062 | 0.4498241867 | -3.382997417 | 0.0007169930654 | 0.03740787103 | FRMD6 |
| ENSG00000081913 | 3142.069415 | 1.504936562 | 0.4573629442 | 3.290464566 | 0.001000220982 | 0.04683851722 | PHLPP1 |
| ENSG00000135097 | 8037.128815 | 1.624869972 | 0.4862571281 | 3.341585919 | 0.0008330123069 | 0.04133730326 | MSI1 |
| ENSG00000121966 | 646.2223644 | 1.633790716 | 0.4857192515 | 3.363652379 | 0.0007691836075 | 0.03899355786 | CXCR4 |
| ENSG00000130475 | 500.2978578 | 1.708429646 | 0.5150307305 | 3.317141181 | 0.0009094364866 | 0.04419839919 | FCHO1 |
| ENSG00000182175 | 3065.498257 | 1.777959459 | 0.4678171152 | 3.800543847 | 0.0001443788667 | 0.01194106607 | RGMA |
| ENSG00000106537 | 569.6948589 | 1.788069506 | 0.5416092211 | 3.301401521 | 0.0009620310034 | 0.04564812493 | TSPAN13 |
| ENSG00000234616 | 915.5773185 | 1.805810209 | 0.5116218306 | 3.529580056 | 0.0004162197571 | 0.02504913282 | JRK |
| ENSG00000137285 | 1549.13591 | 1.807700572 | 0.4570848387 | 3.954846931 | 7.66E-05 | 0.007578136239 | TUBB2B |
| ENSG00000187244 | 830.8973502 | 1.835627555 | 0.529944234 | 3.463812675 | 0.000532577244 | 0.03021694233 | BCAM |
| ENSG00000158246 | 329.1277943 | 1.84030612 | 0.5138695162 | 3.581271242 | 0.0003419264441 | 0.02203172561 | TENT5B |
| ENSG00000169862 | 2213.675792 | 1.844015152 | 0.5178116118 | 3.56116995 | 0.000369206002 | 0.02324879006 | CTNND2 |
| ENSG00000167680 | 862.3468438 | 1.849854275 | 0.5174844977 | 3.574704716 | 0.0003506233826 | 0.02233495886 | SEMA6B |
| ENSG00000196220 | 2807.845705 | 1.85945535 | 0.5061276948 | 3.673885798 | 0.0002388895364 | 0.01693953451 | SRGAP3 |
| ENSG00000183248 | 1923.061861 | 1.874129059 | 0.4871357544 | 3.847241846 | 0.0001194550391 | 0.01039696634 | PRR36 |
| ENSG00000148082 | 1538.462082 | 1.909326231 | 0.4745371918 | 4.023554454 | 5.73E-05 | 0.005956204834 | SHC3 |
| ENSG00000091129 | 1469.280252 | 1.909482935 | 0.5810200104 | 3.286432311 | 0.001014651566 | 0.04709678546 | NRCAM |
| ENSG00000116717 | 534.5990397 | 1.916873031 | 0.5263482575 | 3.641834098 | 0.0002707025137 | 0.01851917114 | GADD45A |

| | | | | | | | |
|---|---|---|---|---|---|---|---|
| ENSG00000111913 | 1603.531023 | 1.917781793 | 0.5026081812 | 3.815659722 | 0.0001358195669 | 0.01134605267 | RIPOR2 |
| ENSG00000175175 | 780.6762074 | 1.930472732 | 0.5228640418 | 3.692112247 | 0.0002223992632 | 0.01623680533 | PPM1E |
| ENSG00000185818 | 641.8307965 | 1.960732991 | 0.5066676373 | 3.869860332 | 0.0001088977211 | 0.009899762957 | NAT8L |
| ENSG00000103460 | 1855.238418 | 1.967071876 | 0.5059629915 | 3.887778175 | 0.0001011660162 | 0.00950160369 | TOX3 |
| ENSG00000150625 | 1685.94854 | 1.982764033 | 0.5380922295 | 3.684803319 | 0.0002288793131 | 0.01640038664 | GPM6A |
| ENSG00000111674 | 1167.978748 | 2.014563255 | 0.4823282803 | 4.176747119 | 2.96E-05 | 0.003698689926 | ENO2 |
| ENSG00000147642 | 232.8135899 | 2.058454601 | 0.5547910521 | 3.710324082 | 0.0002069940936 | 0.01550031447 | SYBU |
| ENSG00000101255 | 862.3294766 | 2.080037714 | 0.5095606911 | 4.082021533 | 4.46E-05 | 0.004974338738 | TRIB3 |
| ENSG00000185274 | 1114.089807 | 2.104398568 | 0.5933333298 | 3.546739181 | 0.0003900305192 | 0.02389445709 | GALNT17 |
| ENSG00000233639 | 1243.890577 | 2.107200685 | 0.5445413543 | 3.869679811 | 0.0001089783819 | 0.009899762957 | PANTR1 |
| ENSG00000167889 | 358.7426521 | 2.111699558 | 0.6226884887 | 3.391261596 | 0.0006957166133 | 0.03683309866 | MGAT5B |
| ENSG00000167614 | 3514.607559 | 2.118955689 | 0.4534482143 | 4.672982763 | 2.97E-06 | 0.0005832726157 | TTYH1 |
| ENSG00000114656 | 354.1183448 | 2.146611962 | 0.6422086537 | 3.342545992 | 0.0008301359726 | 0.04131790541 | CFAP92 |
| ENSG00000007516 | 995.7378283 | 2.167193966 | 0.5908854847 | 3.66770554 | 0.0002447367976 | 0.01709455682 | BAIAP3 |
| ENSG00000040608 | 327.5866096 | 2.211876669 | 0.5572760382 | 3.969086266 | 7.21E-05 | 0.007313420715 | RTN4R |
| ENSG00000175093 | 836.6794258 | 2.242859147 | 0.5662610662 | 3.960821749 | 7.47E-05 | 0.007480026443 | SPSB4 |
| ENSG00000163884 | 174.0398419 | 2.24363629 | 0.6473121511 | 3.46608091 | 0.0005281043303 | 0.0300657753 | KLF15 |
| ENSG00000198822 | 520.7532503 | 2.256255798 | 0.6758927939 | 3.338185905 | 0.0008432730908 | 0.04172194006 | GRM3 |
| ENSG00000155511 | 1029.08815 | 2.256824253 | 0.667927678 | 3.378845236 | 0.0007279097332 | 0.03754454534 | GRIA1 |
| ENSG00000176771 | 1141.84308 | 2.281530612 | 0.6089567512 | 3.746621755 | 0.0001792320287 | 0.01385838719 | NCKAP5 |
| ENSG00000186493 | 294.5770021 | 2.295983918 | 0.6604828438 | 3.476220374 | 0.0005085342287 | 0.02930927831 | IRX2-DT |
| ENSG00000143126 | 9299.719482 | 2.29889342 | 0.5159132639 | 4.455968825 | 8.35E-06 | 0.001361133497 | CELSR2 |
| ENSG00000234745 | 468.2571042 | 2.301406196 | 0.6671217202 | 3.44975456 | 0.0005610964409 | 0.03088631534 | HLA-B |
| ENSG00000065923 | 500.8185616 | 2.309836662 | 0.5972239314 | 3.867622412 | 0.0001099016627 | 0.00992937631 | SLC9A7 |
| ENSG00000178445 | 628.901882 | 2.327615055 | 0.619201909 | 3.759056652 | 0.0001705552191 | 0.01356607638 | GLDC |
| ENSG00000242808 | 809.5543948 | 2.334975744 | 0.692410355 | 3.372242669 | 0.0007455872354 | 0.03813736062 | SOX2-OT |
| ENSG00000168243 | 860.2431555 | 2.346701243 | 0.6880147134 | 3.410830027 | 0.0006476545193 | 0.03507038674 | GNG4 |
| ENSG00000188897 | 209.1160247 | 2.349548511 | 0.6583284089 | 3.568961144 | 0.0003583995696 | 0.02265412336 | |
| ENSG00000183150 | 152.139278 | 2.366332956 | 0.6210414331 | 3.810265837 | 0.0001388174421 | 0.01153850579 | GPR19 |
| ENSG00000081479 | 3128.371394 | 2.367904421 | 0.6268881729 | 3.777235754 | 0.0001585786101 | 0.01279713988 | LRP2 |
| ENSG00000082684 | 3446.39707 | 2.372691032 | 0.6573465979 | 3.609497698 | 0.0003067904958 | 0.02031906455 | SEMA5B |
| ENSG00000109339 | 4045.165195 | 2.388447743 | 0.5777962759 | 4.133719517 | 3.57E-05 | 0.004267932261 | MAPK10 |
| ENSG00000198435 | 677.0598001 | 2.407634165 | 0.5052891341 | 4.764864318 | 1.89E-06 | 0.0003927020465 | NRARP |
| ENSG00000162692 | 1568.471841 | 2.413998859 | 0.5723303826 | 4.217841535 | 2.47E-05 | 0.003178561062 | VCAM1 |

| | | | | | | | |
|---|---|---|---|---|---|---|---|
| ENSG00000172461 | 330.0397313 | 2.422980416 | 0.5416925456 | 4.472980911 | 7.71E-06 | 0.001282319936 | FUT9 |
| ENSG00000170561 | 2298.528109 | 2.425605964 | 0.6493850235 | 3.735235455 | 0.000187539636 | 0.01443360606 | IRX2 |
| ENSG00000117245 | 142.7463482 | 2.430061519 | 0.6201052933 | 3.918788543 | 8.90E-05 | 0.008405996128 | KIF17 |
| ENSG00000161513 | 317.9143117 | 2.436622362 | 0.6787952563 | 3.589627858 | 0.0003311503762 | 0.0214204041 | FDXR |
| ENSG00000214872 | 132.1832293 | 2.4696647 | 0.6792402346 | 3.635922276 | 0.0002769878447 | 0.01871807289 | SMTNL1 |
| ENSG00000134323 | 683.939415 | 2.505028526 | 0.6166344986 | 4.062420334 | 4.86E-05 | 0.005242661736 | MYCN |
| ENSG00000178233 | 534.9409746 | 2.517129911 | 0.6491528445 | 3.877561243 | 0.0001055087732 | 0.009744321367 | TMEM151B |
| ENSG00000129682 | 574.8636143 | 2.543462411 | 0.7273275846 | 3.496997041 | 0.0004705270515 | 0.02734979617 | FGF13 |
| ENSG00000173805 | 154.0245058 | 2.562256341 | 0.6134806035 | 4.176589 | 2.96E-05 | 0.003698689926 | HAP1 |
| ENSG00000116396 | 296.4370628 | 2.582008843 | 0.5375948968 | 4.802889422 | 1.56E-06 | 0.0003466486619 | KCNC4 |
| ENSG00000214357 | 768.0750958 | 2.621029531 | 0.5112670786 | 5.126536873 | 2.95E-07 | 8.47E-05 | NEURL1B |
| ENSG00000106538 | 252.9349839 | 2.628135272 | 0.7350396075 | 3.575501572 | 0.0003495570758 | 0.02233495886 | RARRES2 |
| ENSG00000120756 | 108.0066586 | 2.630312711 | 0.7783173394 | 3.379486204 | 0.0007262145196 | 0.03754454534 | PLS1 |
| ENSG00000233695 | 104.367227 | 2.63202244 | 0.8016512839 | 3.283251076 | 0.001026172263 | 0.04725509057 | GAS6-AS1 |
| ENSG00000155093 | 1048.716996 | 2.652504513 | 0.4902116841 | 5.410936946 | 6.27E-08 | 2.48E-05 | PTPRN2 |
| ENSG00000171766 | 516.5409342 | 2.694120667 | 0.7952846885 | 3.387617926 | 0.0007050240183 | 0.03720736279 | GATM |
| ENSG00000138135 | 133.2818348 | 2.701490144 | 0.65377761 | 4.132123986 | 3.59E-05 | 0.004267932261 | CH25H |
| ENSG00000170837 | 118.0929065 | 2.72013706 | 0.666264272 | 4.082669856 | 4.45E-05 | 0.004974338738 | GPR27 |
| ENSG00000188290 | 421.2161192 | 2.731697453 | 0.7697709804 | 3.548714518 | 0.0003871165331 | 0.02383490832 | HES4 |
| ENSG00000137642 | 1164.466562 | 2.738831042 | 0.7661762292 | 3.574675039 | 0.0003506631534 | 0.02233495886 | SORL1 |
| ENSG00000168918 | 312.6510102 | 2.747625513 | 0.7859288191 | 3.496023362 | 0.000472247351 | 0.02735414621 | INPP5D |
| ENSG00000171724 | 1812.66575 | 2.748884163 | 0.4912675338 | 5.595493238 | 2.20E-08 | 9.91E-06 | VAT1L |
| ENSG00000127578 | 282.2604604 | 2.751598817 | 0.6885438126 | 3.996258141 | 6.44E-05 | 0.006603582484 | WFIKKN1 |
| ENSG00000029534 | 141.4873128 | 2.768409255 | 0.6304879406 | 4.390899614 | 1.13E-05 | 0.001737556647 | ANK1 |
| ENSG00000259905 | 141.7607546 | 2.773796503 | 0.7902610412 | 3.509975007 | 0.0004481488161 | 0.02632517992 | PWRN1 |
| ENSG00000124191 | 347.062202 | 2.794418873 | 0.6541235485 | 4.272004699 | 1.94E-05 | 0.00270626836 | TOX2 |
| ENSG00000182968 | 2971.503922 | 2.812342597 | 0.4923999826 | 5.711500196 | 1.12E-08 | 5.48E-06 | SOX1 |
| ENSG00000174498 | 4390.845523 | 2.845430167 | 0.8114819181 | 3.506461578 | 0.0004541069919 | 0.02658124871 | IGDCC3 |
| ENSG00000130054 | 142.3830563 | 2.863011361 | 0.7668523499 | 3.733458418 | 0.0001888683873 | 0.01446018174 | NALF2 |
| ENSG00000143195 | 3114.535972 | 2.872977747 | 0.5912651868 | 4.859034172 | 1.18E-06 | 0.0002801376136 | ILDR2 |
| ENSG00000139116 | 1432.49956 | 2.884044572 | 0.7961848361 | 3.622330446 | 0.0002919608272 | 0.01941422716 | KIF21A |
| ENSG00000148204 | 5157.123675 | 2.945730847 | 0.5338639528 | 5.517755659 | 3.43E-08 | 1.44E-05 | CRB2 |
| ENSG00000129159 | 351.8015863 | 2.995618404 | 0.8259646401 | 3.626811923 | 0.0002869421391 | 0.01915713301 | KCNC1 |
| ENSG00000110042 | 2755.640885 | 3.00409717 | 0.4819527178 | 6.233178192 | 4.57E-10 | 3.45E-07 | DTX4 |

| | | | | | | | |
|---|---|---|---|---|---|---|---|
| ENSG00000176244 | 588.8363004 | 3.014660561 | 0.694339962 | 4.341764446 | 1.41E-05 | 0.002079368508 | ACBD7 |
| ENSG00000082556 | 98.27307843 | 3.026598736 | 0.8071740875 | 3.749623264 | 0.0001771004294 | 0.0138407525 | OPRK1 |
| ENSG00000175161 | 655.4802685 | 3.028159993 | 0.6891931702 | 4.393775393 | 1.11E-05 | 0.001730743476 | CADM2 |
| ENSG00000204128 | 89.65720599 | 3.033487257 | 0.7130839419 | 4.254039501 | 2.10E-05 | 0.002814660922 | C2orf72 |
| ENSG00000188064 | 306.2802649 | 3.035675043 | 0.8099213216 | 3.748111035 | 0.0001781713808 | 0.0138407525 | WNT7B |
| ENSG00000251141 | 92.94096097 | 3.035702568 | 0.8172287641 | 3.714630103 | 0.0002035011495 | 0.01530770638 | MRPS30-DT |
| ENSG00000184261 | 184.2561688 | 3.054557114 | 0.7292909874 | 4.188392791 | 2.81E-05 | 0.003565111597 | KCNK12 |
| ENSG00000106278 | 13907.20391 | 3.070999896 | 0.8167210099 | 3.760157825 | 0.0001698061936 | 0.01356607638 | PTPRZ1 |
| ENSG00000109255 | 89.3621255 | 3.088187963 | 0.7819884534 | 3.949147778 | 7.84E-05 | 0.007594696 | NMU |
| ENSG00000198576 | 117.3987254 | 3.088751887 | 0.9230224663 | 3.346345295 | 0.0008188435798 | 0.04108048187 | ARC |
| ENSG00000150394 | 361.5359953 | 3.094009973 | 0.8963411354 | 3.451821914 | 0.0005568151032 | 0.03083915752 | CDH8 |
| ENSG00000164076 | 212.2150783 | 3.107989922 | 0.5911809301 | 5.257256727 | 1.46E-07 | 4.67E-05 | CAMKV |
| ENSG00000145423 | 2162.400928 | 3.132721335 | 0.9588464033 | 3.267177438 | 0.001086255744 | 0.04960965795 | SFRP2 |
| ENSG00000231566 | 91.75478247 | 3.162015234 | 0.7974493188 | 3.965161371 | 7.33E-05 | 0.007389751864 | LINC02595 |
| ENSG00000224945 | 198.7425184 | 3.174722678 | 0.8317599106 | 3.816873881 | 0.0001351532102 | 0.01134605267 | |
| ENSG00000105855 | 8594.497329 | 3.245378304 | 0.7955840618 | 4.07923997 | 4.52E-05 | 0.004974338738 | ITGB8 |
| ENSG00000122733 | 52.53509629 | 3.246730107 | 0.9419550971 | 3.446799234 | 0.0005672699661 | 0.03102071025 | PHF24 |
| ENSG00000141449 | 1546.286596 | 3.246957045 | 0.7027873735 | 4.620112949 | 3.84E-06 | 0.0007245252932 | GREB1L |
| ENSG00000127241 | 213.0458894 | 3.254803044 | 0.6126557494 | 5.312613236 | 1.08E-07 | 3.74E-05 | MASP1 |
| ENSG00000080573 | 1537.032323 | 3.255480937 | 0.4728303262 | 6.885093355 | 5.77E-12 | 8.73E-09 | COL5A3 |
| ENSG00000197629 | 52.61157599 | 3.268556247 | 0.9239559497 | 3.537567184 | 0.0004038314329 | 0.02459059627 | MPEG1 |
| ENSG00000171956 | 675.7121664 | 3.268807442 | 0.7963009644 | 4.104989933 | 4.04E-05 | 0.004667793948 | FOXB1 |
| ENSG00000189410 | 116.68116 | 3.27447054 | 0.7952757936 | 4.117402499 | 3.83E-05 | 0.004485746957 | SH2D5 |
| ENSG00000112232 | 130.5769555 | 3.282939528 | 0.6764252684 | 4.853366191 | 1.21E-06 | 0.0002842077757 | KHDRBS2 |
| ENSG00000163661 | 1858.456359 | 3.319736549 | 0.7414498243 | 4.477358332 | 7.56E-06 | 0.001269004693 | PTX3 |
| ENSG00000258708 | 41.58015823 | 3.325370293 | 0.9768378172 | 3.404219446 | 0.0006635342326 | 0.0358136139 | SLC25A21-AS1 |
| ENSG00000170500 | 2273.736926 | 3.335113773 | 0.724597467 | 4.602712437 | 4.17E-06 | 0.0007789443067 | LONRF2 |
| ENSG00000166450 | 13569.9426 | 3.336949343 | 0.9877738217 | 3.37825246 | 0.0007294807595 | 0.03754454534 | PRTG |
| ENSG00000162551 | 175.3245955 | 3.351999628 | 0.6302029281 | 5.318921063 | 1.04E-07 | 3.74E-05 | ALPL |
| ENSG00000255545 | 115.01926 | 3.363562069 | 0.9052018297 | 3.715814483 | 0.0002025501619 | 0.01530542678 | B3GAT1-DT |
| ENSG00000070808 | 214.9882116 | 3.414681924 | 0.6344914786 | 5.381761676 | 7.38E-08 | 2.79E-05 | CAMK2A |
| ENSG00000164920 | 50.71771511 | 3.481720461 | 0.9288376734 | 3.748470331 | 0.0001779163794 | 0.0138407525 | OSR2 |
| ENSG00000136014 | 156.2814501 | 3.498014591 | 0.7785241104 | 4.493135851 | 7.02E-06 | 0.001202788927 | USP44 |
| ENSG00000222012 | 214.2538053 | 3.502334513 | 0.6498926054 | 5.389097343 | 7.08E-08 | 2.74E-05 | |

| | | | | | | | |
|---|---|---|---|---|---|---|---|
| ENSG00000140067 | 190.5025152 | 3.508370018 | 0.7820182045 | 4.486302234 | 7.25E-06 | 0.001229326509 | FAM181A |
| ENSG00000206557 | 2060.781656 | 3.524251069 | 0.6086393547 | 5.790376587 | 7.02E-09 | 3.54E-06 | TRIM71 |
| ENSG00000127084 | 50.61926406 | 3.526716734 | 1.000184344 | 3.526066724 | 0.0004217806894 | 0.02511986917 | FGD3 |
| ENSG00000185559 | 3405.409797 | 3.537907582 | 0.9499723681 | 3.724221568 | 0.000195918805 | 0.01487193705 | DLK1 |
| ENSG00000243742 | 192.6517636 | 3.613480742 | 0.9467993807 | 3.81652208 | 0.0001353459684 | 0.01134605267 | RPLP0P2 |
| ENSG00000197921 | 224.5823076 | 3.748160983 | 1.084562299 | 3.455920408 | 0.0005484172312 | 0.03073629702 | HES5 |
| ENSG00000137843 | 94.16978098 | 3.764646452 | 1.060600017 | 3.549544023 | 0.0003858989354 | 0.02383490832 | PAK6 |
| ENSG00000109906 | 798.4564627 | 3.794966394 | 0.7924816499 | 4.788712009 | 1.68E-06 | 0.0003671610906 | ZBTB16 |
| ENSG00000250366 | 284.1021432 | 3.813573767 | 0.5525202199 | 6.902143359 | 5.12E-12 | 8.52E-09 | TUNAR |
| ENSG00000069431 | 534.2817666 | 3.813824608 | 0.874863838 | 4.359335067 | 1.30E-05 | 0.00196739385 | ABCC9 |
| ENSG00000258498 | 374.5982868 | 3.832993655 | 1.129863501 | 3.392439576 | 0.0006927320896 | 0.03683309866 | DIO3OS |
| ENSG00000148600 | 106.9298213 | 3.842180297 | 1.013071644 | 3.792604718 | 0.0001490753529 | 0.01220802299 | CDHR1 |
| ENSG00000139352 | 335.0759754 | 3.845907895 | 0.8066758017 | 4.767600425 | 1.86E-06 | 0.0003923118216 | ASCL1 |
| ENSG00000234690 | 92.29774546 | 3.848743515 | 0.7241590431 | 5.314776569 | 1.07E-07 | 3.74E-05 | EPCAM-DT |
| ENSG00000153714 | 132.3372381 | 3.878326806 | 0.9832526251 | 3.944384899 | 8.00E-05 | 0.007687885952 | LURAP1L |
| ENSG00000138823 | 339.2335714 | 3.903718827 | 0.9501329964 | 4.108602524 | 3.98E-05 | 0.00462752277 | MTTP |
| ENSG00000210140 | 69.57742106 | 3.904640514 | 1.162481435 | 3.358884192 | 0.0007825785555 | 0.03954281431 | MT-TC |
| ENSG00000049130 | 1790.979964 | 3.921812824 | 0.9970901533 | 3.933258002 | 8.38E-05 | 0.008006481874 | KITLG |
| ENSG00000104267 | 83.1071795 | 3.955570839 | 1.054894152 | 3.749732457 | 0.0001770233344 | 0.0138407525 | CA2 |
| ENSG00000207955 | 162.8713637 | 3.974771081 | 1.04082492 | 3.818866175 | 0.0001340664648 | 0.01134605267 | MIR219A2HG |
| ENSG00000210196 | 1760.502475 | 4.133877875 | 1.168883469 | 3.536603934 | 0.0004053069886 | 0.02459059627 | MT-TP |
| ENSG00000187122 | 2132.749658 | 4.169579866 | 1.02917332 | 4.051387444 | 5.09E-05 | 0.005425691825 | SLIT1 |
| ENSG00000250337 | 2011.9105 | 4.217646996 | 1.196413644 | 3.525241472 | 0.0004230969303 | 0.02511986917 | PURPL |
| ENSG00000117600 | 165.1683621 | 4.220457738 | 0.7284330557 | 5.793885526 | 6.88E-09 | 3.54E-06 | PLPPR4 |
| ENSG00000174721 | 3513.119219 | 4.226562895 | 0.6491245092 | 6.511174412 | 7.46E-11 | 7.75E-08 | FGFBP3 |
| ENSG00000258548 | 174.8423169 | 4.302524579 | 1.285048336 | 3.348142212 | 0.0008135525261 | 0.04098332483 | LINC00645 |
| ENSG00000057657 | 341.8220987 | 4.395732934 | 0.8538062582 | 5.148396245 | 2.63E-07 | 8.09E-05 | PRDM1 |
| ENSG00000224243 | 232.3608344 | 4.526173304 | 0.8600968868 | 5.262399357 | 1.42E-07 | 4.63E-05 | SOX1-OT |
| ENSG00000154764 | 170.6118311 | 4.544638855 | 0.9319952328 | 4.87624689 | 1.08E-06 | 0.0002643295904 | WNT7A |
| ENSG00000173452 | 65.13708665 | 4.593432638 | 1.193850926 | 3.847576393 | 0.0001192920857 | 0.01039696634 | TMEM196 |
| ENSG00000158769 | 101.4332563 | 4.643900468 | 1.224304517 | 3.793092653 | 0.0001487826096 | 0.01220802299 | F11R |
| ENSG00000229520 | 245.521282 | 4.799570767 | 0.6390485989 | 7.510494156 | 5.89E-14 | 1.40E-10 | LINC00404 |
| ENSG00000066735 | 439.9194781 | 4.894512283 | 0.6653724809 | 7.356048564 | 1.89E-13 | 3.50E-10 | KIF26A |
| ENSG00000227640 | 589.7296711 | 4.952111943 | 0.6476027266 | 7.646836154 | 2.06E-14 | 5.71E-11 | SOX21-AS1 |

| | | | | | | | |
|---|---|---|---|---|---|---|---|
| ENSG00000105825 | 2254.548928 | 4.975018138 | 1.231162301 | 4.040911693 | 5.32E-05 | 0.005602057192 | TFPI2 |
| ENSG00000004848 | 102.9662691 | 5.109714507 | 0.8690572819 | 5.879606113 | 4.11E-09 | 2.28E-06 | ARX |
| ENSG00000181965 | 116.6963242 | 5.163661359 | 1.405886359 | 3.672886736 | 0.0002398258007 | 0.01693953451 | NEUROG1 |
| ENSG00000138083 | 159.6057976 | 5.189351761 | 1.216149876 | 4.267033087 | 1.98E-05 | 0.002744203879 | SIX3 |
| ENSG00000238230 | 89.62932593 | 5.256383793 | 1.14474452 | 4.591752747 | 4.40E-06 | 0.000811877217 | LINC00391 |
| ENSG00000131914 | 562.9934588 | 5.289018326 | 1.038173739 | 5.094540658 | 3.50E-07 | 9.69E-05 | LIN28A |
| ENSG00000125285 | 1463.130576 | 5.39258163 | 0.6049788378 | 8.913669856 | 4.94E-19 | 2.60E-15 | SOX21 |
| ENSG00000198732 | 1002.296528 | 5.39363868 | 0.860918048 | 6.264985027 | 3.73E-10 | 2.95E-07 | SMOC1 |
| ENSG00000168348 | 179.5508954 | 5.530329497 | 1.425045842 | 3.880808136 | 0.0001041099494 | 0.009723167408 | INSM2 |
| ENSG00000260343 | 116.2342986 | 5.577778126 | 1.102097373 | 5.061057454 | 4.17E-07 | 0.0001136257327 | LINC01043 |
| ENSG00000169627 | 76.65925214 | 5.72407919 | 1.499878531 | 3.81636184 | 0.0001354338532 | 0.01134605267 | BOLA2B |
| ENSG00000231827 | 399.7640027 | 5.970686802 | 0.9632916684 | 6.198212855 | 5.71E-10 | 3.96E-07 | |
| ENSG00000264462 | 73.38850443 | 5.981898592 | 1.624196706 | 3.682988994 | 0.0002305151154 | 0.01644670935 | MIR3648-2 |
| ENSG00000176956 | 38.01536868 | 6.319286363 | 1.460762935 | 4.326017735 | 1.52E-05 | 0.002214042709 | LY6H |
| ENSG00000014257 | 148.9221963 | 6.482570398 | 0.8751904067 | 7.407040055 | 1.29E-13 | 2.68E-10 | ACP3 |
| ENSG00000130600 | 543.0571653 | 6.523594641 | 0.9871030507 | 6.608828366 | 3.87E-11 | 4.29E-08 | H19 |
